# Sse1, Hsp110 chaperone of yeast, controls the cellular fate during Endoplasmic Reticulum-stress

**DOI:** 10.1101/2023.06.29.547006

**Authors:** Mainak Pratim Jha, Vignesh Kumar, Lakshita Sharma, Asmita Ghosh, Koyeli Mapa

## Abstract

Sse1 is a cytosolic Hsp110 molecular chaperone of yeast, *Saccharomyces cerevisiae*. Its multifaceted roles in cellular protein homeostasis as Nucleotide Exchange Factor (NEF), as protein-disaggregase and as a Chaperone linked to Protein Synthesis (CLIPS), are well documented. In the currently study, we show that *SSE1* genetically interacts with *IRE1* and *HAC1*, the Endoplasmic Reticulum-Unfolded Protein Response (ER-UPR) sensors implicating its role in ER protein homeostasis. Interestingly, absence of this chaperone imparts unusual resistance to tunicamycin-induced ER stress which depends on the intact Ire1-Hac1 mediated ER-UPR signalling. Furthermore, cells lacking *SSE1* show ER-stress-responsive inefficient reorganization of translating ribosomes from polysomes to monosomes and increased monosome content that drive uninterrupted protein translation. In consequence, the kinetics of ER-UPR is starkly different in *sse1*Δ strain where we show that stress response induction and restoration of homeostasis is prominently faster in contrast to the wildtype (WT) cells. Importantly, Sse1 plays a critical role in controlling the ER-stress mediated cell division arrest which is escaped in *sse1*Δ strain during chronic tunicamycin stress. Consequently, *sse1*Δ strain shows significantly higher cell viability in comparison to WT yeast, following short-term as well as long-term tunicamycin stress. In summary, we demonstrate a new role of Sse1 in ER protein homeostasis where the chaperone genetically interacts with ER-UPR pathway, controls the protein translation during ER stress and the kinetics of ER-UPR. More importantly, we show the crtical role of Sse1 in regulating the ER-stress-induced cell division arrest and cell death during global ER stress by tunicamycin.

**Author Summary:** Sse1 is a cytosolic Hsp110 molecular chaperone of yeast, *Saccharomyces cerevisiae*. It performs various functions as Nucleotide Exchange Factor (NEF), as a protein-disaggregase and as a Chaperone linked to Protein Synthesis (CLIPS) to maintain a healthy proteome in the cytosol. In the present study, we show that *SSE1* is a critical player to maintain Endoplasmic Reticulum (ER) protein homeostasis. We report that *SSE1* genetically interacts with the ER-Unfolded Protein Response (ER-UPR) sensors, *IRE1* and *HAC1*. In absence of *SSE1,* yeast cells gain unusual fitness to tunicamycin (Tm)-induced ER stress which depends on the functional Ire1-Hac1 mediated ER-UPR signalling. Furthermore, Sse1 controls the ribosomal organization during tunicamycin-induced ER-stress and in absence of the chaperone the amount of polysomes and monosmes are high leading to uninterrupted protein translation during ER stress. Consequently, *sse1*Δ strain exhibits starkly different ER-UPR kinetics compared to wildtype (WT) cells. Importantly, our data reveal that cell division arrest and cell death due to Tm-induced ER-stress is critically controlled by Sse1.

## Introduction

Sse1 is a member of the Hsp110 group of molecular chaperones found in yeast, *Saccharomyces cerevisiae*. To date, it is well established that Hsp110s act as nucleotide exchange factors (NEF) for its Hsp70 partners [1–4]. Hsp110s are prominently similar to Hsp70 chaperones in their domain organization and structure [4–6]. Like Hsp70s, Hsp110s are also two-domain proteins, with a ~45 kDa N-terminal Nucleotide Binding Domain (NBD) and a ~25 kDa Peptide or Substrate Binding Domain (PBD or SBD) at the C-terminus. The structural and molecular basis of the NEF function of yeast Hsp110, Sse1, has been explored in great detail in the last decade [1, 4, 6]. Although there is plethora of information regarding domain allostery of Hsp70s, the same is still elusive for Sse1 and Hsp110s, in general. Recently, we showed that the ATP-hydrolysis-driven conventional domain movements found in Hsp70s are absent in Sse1, although there are distinctive non-canonical domain movements found in this protein upon ATP hydrolysis [7]. Sse1 is also distinctly different from its Hsp70 partners in its cellular functions. Apart from its well-known function as a NEF for Hsp70 partner proteins like Ssa1 or Ssb1 [3, 8], Sse1 is also known to act as a co-chaperone for Hsp90 [9] or assists in protein disaggregation in association with Hsp70 and Hsp104 chaperones [7]. Due to its co-regulated expression pattern with cytosolic translation machinery, Sse1 is additionally categorized as a <u>C</u>haperones <u>L</u>inked to <u>P</u>rotein <u>S</u>ynthesis (CLIPS) [10]. CLIPS are a group of cytosolic chaperones that are transcriptionally downregulated during stress (under different cellular stresses like heat shock, osmotic shock, oxidative stress etc.) and are transcriptionally co-regulated with protein translation apparatus. They are distinct from the stress-induced chaperones designated commonly as Heat Shock Proteins or HSPs. Among CLIPS, only Ssa1 and Sse1 were found to be overexpressed during heat stress but were transcriptionally repressed during most of the other stresses [10]. Very recently, another function of Sse1 in cellular protein homeostasis was revealed. It was shown that cells deleted of *SSE1* are less efficient in ER (Endoplasmic Reticulum)-reflux of proteins, a newly described quality control pathway parallel to ERAD (ER-<u>A</u>ssociated <u>D</u>egradation of proteins) which is important for upkeep of ER protein homeostasis [11]. Earlier, it was also shown that some of the ERAD substrates are stabilized by Sse1 [12]. Thus, it is evident that Sse1 has a cross-compartment role in maintaining protein homeostasis in yeast.

In this work, in an attempt to understand the breadth of Sse1’s role in cellular proteostasis, we subjected the *SSE1* deletion strain to known proteotoxic stressors. To most of the tested stressors, *sse1*Δ strain does not exhibit any apparent hypersensitivity except for protein translation blockers like cycloheximide. Interestingly, when treated with ER stressor tunicamycin (inhibits N-linked glycosylation thereby preventing correct folding of N-linked glycosylated proteins leading to ER stress), *sse1*Δ strain shows significant fitness indicating the important regulatory role of Sse1 during ER stress. We further found that the tunicamycin resistance of *sse1*Δ strain depends on the activation of Ire1-Hac1 mediated ER-UPR (unfolded Protein Response) signalling. Notably, *SSE1* shows negative genetic interaction with *IRE1* and *HAC1* even in the absence of ER stress further associating its functions to basal ER-UPR. Moreover, we show that in the absence of *SSE1*, ER-stress-induced changes in ribosome organization are different. The extent of polysome to monosome transition following Tm-induced ER stress is far less prominent in *sse1*Δ cells compared to WT cells. These inefficient alterations in ribosome organization lead to altered kinetics of synthesis of UPR-induced proteins in *sse1*Δ strain where we show that the response to ER stress is quicker and short-lasting compared to WT cells. Importantly, ER-stress-induced cell division arrest as observed in WT cells is evaded in *sse1*Δ strain during Tm-induced ER stress indicating an important role of Sse1 in suspending the cell division following ER stress. Thus, we show that Sse1 plays a key role in maintaining ER protein homeostasis and stress-induced cell division arrest after Tm-induced ER stress. In conclusion, Sse1 despite being a cytosolic chaperone plays a crucial role in deciding cellular fate during global ER stress induced by tunicamycin.

## Results

### The absence of Sse1 confers tunicamycin resistance to yeast

To understand the possible non-canonical roles of Sse1 and overall Hsp110 molecular chaperones, during proteotoxic stresses of varied origin, we checked the growth phenotype of deletion strain of *SSE1* (*sse1*Δ) during known proteotoxic conditions. We checked the phenotype of the strain during heat stress (37°C), oxidative stress (induced by H_2_O_2_), protein translation block (induced by cycloheximide or by a limited supply of carbon source), ER stress [induced by N-linked glycosylation blocker Tunicamycin (Tm)], general protein misfolding stress induced by AZC (L-Azetidine-2-Carboxylic Acid), Hsp90 inhibition (by geldanamycin), DNA damage and mitochondrial stress by ethidium bromide and by CCCP (mitochondrial oxidative phosphorylation uncoupler) treatment. We kept the wildtype (WT) yeast strain (BY4741) and the deletion strain of *SSE1* paralog, *SSE2,* (*sse2*Δ strain) [13] as controls. The *sse1*Δ strain exhibits a prominent growth phenotype at a permissive temperature (30°C) in the absence of any additional stress in comparison to WT and *sse2*Δ strains (Figure 1Ai, left panel and 1Aii). A similar phenotype of the *sse1*Δ strain is observed during heat shock at 37°C (Figure 1Ai, right panel). Notably, *sse1*Δ strain shows higher sensitivity towards protein translation block by cycloheximide (Figure 1Bi and Bii) while the strain is not overtly sensitive to other stressors (Figure 1Aiii). Surprisingly, in the presence of ER stressor tunicamycin (Tm), *sse1*Δ strain exhibits a significant fitness advantage in comparison to the WT or *sse2*Δ strains (Figure 1Ci and Cii). Induction of ER-UPR by the same amount of Tm (as used for checking the growth phenotype) is confirmed by significant splicing of *HAC1* mRNA (Figure 1Di and Dii) and overexpression of ER-resident Hsp70 chaperone, Kar2 (yeast orthologue of human BiP protein) (Figure 1Ei and Eii). The specificity of the Tm-resistance phenotype of the *sse1*Δ strain is further confirmed by a complementation assay by expressing *SSE1* under its native promoter by a centromeric plasmid in the *sse1*Δ strain (Figure S1A and B). Expression of *SSE1* in *sse1*Δ strain reverted the sensitivity to Tm-induced ER stress like WT yeast cells (Figure S1A, right panel). Importantly, overexpression of *SSE1* successfully restored the Tm-sensitivity similar to the endogenous level of expression although overexpression of another cytosolic NEF, *FES1* could not complement the phenotype (Figure S1A, right panel). This data indicates that although *FES1* overexpression can rescue the synthetic lethal phenotype *sse1*Δ-*sse2*Δ double deletion strain (lacking NEF activity of Hsp110s) as shown previously [7], the role of Sse1 during ER stress cannot be accomplished by Fes1 indicating a possible non-NEF additional function of Sse1 during Tm-induced ER stress. In contrast, overexpression of *SSE2* in *sse1*Δ strain leads to reversal of Tm-sensitivity indicating a Hsp110-specific role during ER stress (Figure S1B, right panel). We reiterate that the physiological level of Sse2 is not sufficient to accomplish Sse1’s function during ER stress and the Tm-sensitivity of *sse1*Δ strain is regained only after *SSE2* overexpression. Moreover, expression of the ATP hydrolysis-deficient mutant of Sse1 (K69Q) in *sse1*Δ cells restored the Tm-sensitivity like WT-Sse1 while the ATP-binding-deficient mutant (Sse1G205D) could not complement the phenotype like WT-Sse1 (Figure S1C, right panel). Another mutant (Sse1G233D) which is reported to be deficient in interaction with Ssa1 and ATP-binding [8, 14], also could not complement the phenotype like WT-Sse1. This data indicates the importance of ATP-binding and not ATP-hydrolysis by Sse1 for its function, during ER stress. Importantly, the *sse1*Δ strain does not exhibit any fitness when treated with subcritical concentrations of Tm (Figure 1F) which is not adequate to mount ER-UPR (Figure 1Gi and Gii and Figure S1D, upper and lower panels). This finding indicates that tunicamycin-resistance of the *sse1*Δ strain is dependent on the efficient mounting of ER-UPR. To check any adaptive changes in the glycosylation status of proteins in *the sse1*Δ strain, we specifically captured the glycosylated proteins using concanavalin A (ConA) from both *sse1*Δ and WT cells. There was no visible change in the glycosylated proteins in the *sse1*Δ strain (Figure S1E) ruling out the possibility of adaptive enhanced glycosylation or proteins in the *sse1*Δ strain which may confer growth fitness during Tm-induced ER stress.

**Figure 1:**
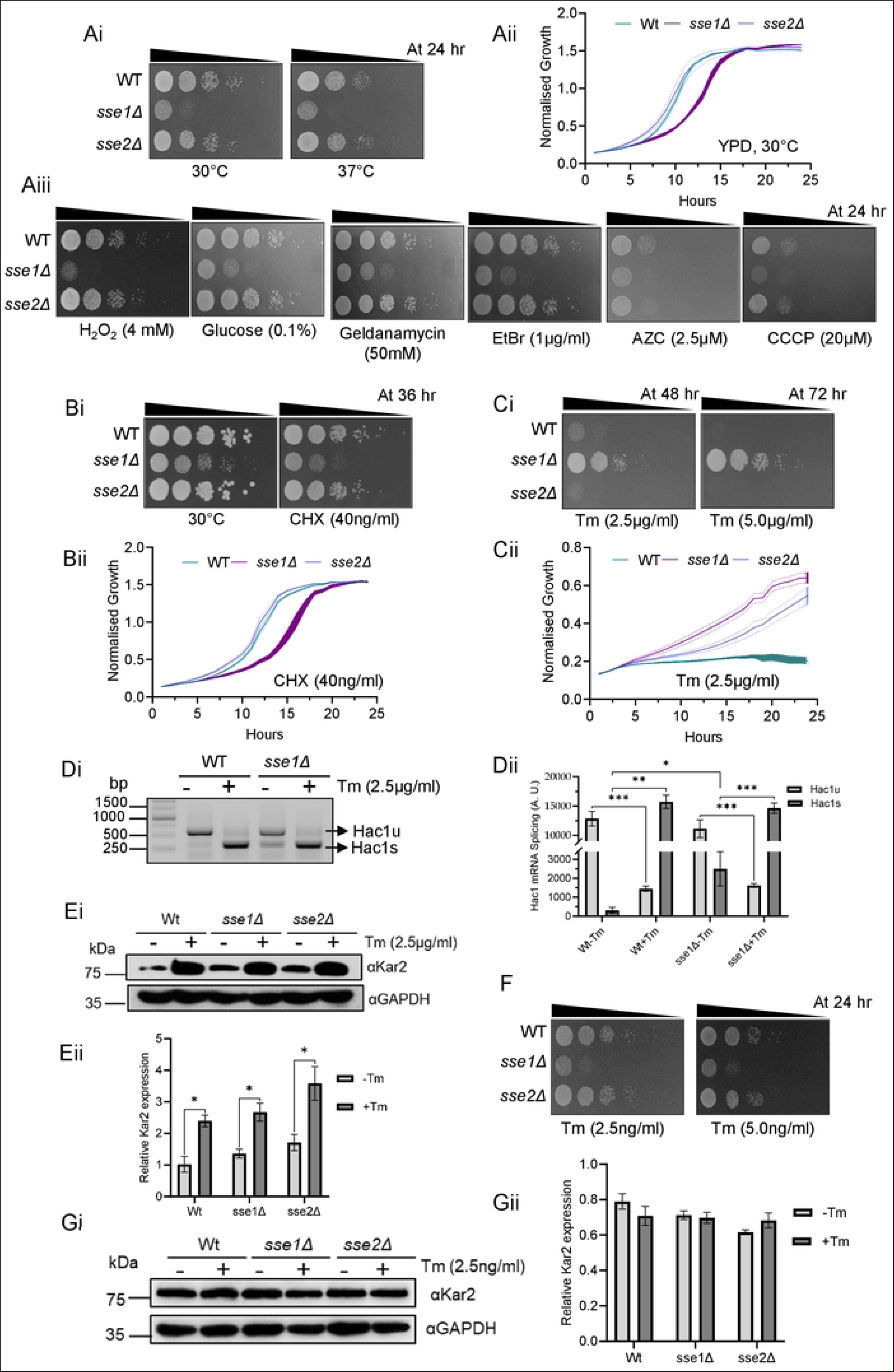
*SSE1* deletion imparts resistance to tunicamycin-induced ER stress in yeast. **A. (Ai)** Yeast growth assay by serial drop dilutions using the strains WT (BY4741), *sse1Δ*, and *sse2Δ* in YPAD plates at permissive temperature (30°C) and under heat stress (37°C) (Ai right panel). **(Aii)** Growth curves in liquid YPD media at permissive temperature (30°C) of the yeast strains used for drop dilution assay in panel Ai. The normalised growth is plotted as line plot for each strain with shaded area representing the error range for the measurements. **(Aiii)** Drop dilution assay with the same strains as in panel Ai in presence of the following proteotoxic stresses: oxidative stress [4mM hydrogen peroxide(H_2_O_2_)], nutrient starvation (0.1% glucose), Hsp90 inhibition by geldanamycin (50mM), DNA damaging agent and mitochondrial stressor (Ethidium Bromide, 1µg/ml), general proteotoxic stress by proline analogue L-Azetidine-2-Carboxylic Acid (AZC, 2.5 µM), and mitochondrial oxidative phosphorylation inhibitor and protonophore, CCCP, (20µM Carbonyl Cyanide m-Chloro-Phenyl hydrazone). All treatments with stressors were done at 30°C. The triangle above each panel indicates the increasing dilutions. The time mentioned in hours represents the time of incubation before taking the image of the plates. Aii. The growth curves of WT, *sse1Δ*, and *sse2Δ* done in liquid media (YPD) at 30°C are shown. **B.** (**Bi)** Drop dilution assay using the WT, *sse1*Δ, and *sse2*Δ strains in YPD plates at permissive temperature (30°C) and in presence of protein translation blocker Cycloheximide (40 ng/ml). (**Bii)** Growth curves of three strains used in panel (Bi) in liquid media in presence of cycloheximide (40 ng/ml) in YPD are shown as stated earlier in (Aii). **C. (Ci)** Similar to panels A and B, a drop dilution assay was done in presence of ER stressor Tunicamycin (Tm); in two concentrations 2.5 µg/ml and 5.0 µg/ml sufficient to elicit ER-UPR. **(Cii)** Growth curves of three strains used in panel (Ci) in liquid media in presence of tunicamycin (2.5 μg/ml) in YPD are shown as stated earlier in (Aii). **D. (Di)** The presence of *HAC1* mRNA splice variants was checked from untreated yeast strains (WT, *sse1*Δ) and the same strains after treatment with tunicamycin (2.5 µg/ml) by synthesizing the cDNAs followed by PCR amplifications with the help of specific primers. **(Dii)** The band intensities were quantified using densitometry and were plotted as a bar plot with whisks representing SEM as shown in the bottom panel (n=3). Statistical significance was calculated using unpaired T-tests and the significant pairs were plotted in the graph (Hac1u-WT-Tm/Hac1u-WT+Tm, p=0.0008, ***; Hac1s-WT-Tm/Hac1s-WT+Tm, p-value=0.0002,***; Hac1u-sse1Δ-Tm/Hac1u-sse1Δ+Tm, p=0.0033, **; and Hac1s-sse1Δ-Tm/Hac1s-sse1Δ+Tm, p=0.0006, ***). All the above pairwise comparisons are significant even after Bonferroni correction with the only exception being Hac1s-WT-Tm/Hac1s-sse1Δ-Tm (one-tailed p=0.0383, *) showed marginal significance and plotted in the graph. **E. (Ei)** Western blot showing the distinct increase in Kar2 (ER-resident Hsp70 and ER-UPR marker) levels in response to ER stress by optimum (2.5 µg/ml) concentration of Tm signifying proper mounting of ER-UPR. GAPDH was used as the loading control. **(Eii)** The bands were quantified by densitometry and were plotted as a bar plot with whisks representing SEM in the bottom panel (n=3). Statistical significance was calculated using unpaired T-tests and the significant pairs were plotted in the graph (WT-Tm/WT+Tm, p=0.0112, *; *sse1*Δ-Tm/*sse1*Δ+Tm, p=0.0142, *; and *sse2*Δ-Tm/*sse2*Δ+Tm, p=0.0347, *). The above pairwise comparisons, except the last pair (*sse2*Δ-Tm/*sse*2Δ+Tm), are significant even after Bonferroni correction. **F.** Drop dilution assay using the same strains as shown in panels A-C, in presence of the ER stressor Tm in sub-optimal (2.5 ng/ml and 5.0 ng/ml) concentrations. **G. (Gi)** Western blot showing no prominent change in Kar2 levels in response to suboptimal (2.5 ng/ml) concentration of Tunicamycin indicating no activation of ER-UPR at this concentration of Tm. GAPDH was used as the loading control. **(Gii)** The bands were quantified by densitometry and were plotted as a bar plot with whisks representing SEM in the right-side panel. Statistical significance was calculated using unpaired T-tests and none of the pairwise comparisons was found to be significant.

### *SSE1* exhibits negative genetic interaction with *IRE1* and *HAC1* and tunicamycin resistance of *sse1*Δ strain depends on the presence of functional ER-UPR signalling by the Ire1-Hac1 pathway

The absence of growth fitness of *sse1*Δ strain at subcritical Tm concentrations hinted towards the necessity of a threshold of ER stress that is sufficient to mount ER-UPR, for gaining cellular fitness against ER stress. As ER-UPR induction is solely dependent on the Ire1-Hac1 pathway in yeast, to understand the Ire1-Hac1 signalling dependence of Tm-resistance of *sse1*Δ strain, we deleted *SSE1* in *ire1*Δ or *hac1*Δ strains to generate the double knockouts of the *ire1*Δ-*sse1*Δ and *hac1*Δ-*sse1*Δ strains. Single deletion strains, *ire1*Δ or *hac1*Δ, grow similarly to WT cells at permissive temperature (30°C) (Figure 2A, left panel) as well as during heat stress (37°C) (Figure 2A, right panel) indicating no alterations in growth rate under physiological conditions or even during heat stress in absence of the ER-UPR sensors. The double knockout strains of *ire1*Δ-*sse1*Δ and *hac1*Δ-*sse1*Δ, show synthetic growth defects at a permissive temperature in the absence of any additional stress indicating a negative genetic interaction between *SSE1* and *IRE1* as well as between *SSE1* and *HAC1* (Figure 2B, left panel). To the best of our knowledge, any experimental evidence of the genetic interaction between *SSE1* and *IRE1-HAC1* pathway is hitherto unknown in literature and this finding implicates an important role of *SSE1* in ER-UPR. During heat stress at 37°C, the synthetic growth sickness of *ire1*Δ-*sse1*Δ and *hac1*Δ-*sse1*Δ strains is significantly aggravated compared to *sse1*Δ strain indicating a stronger genetic interaction with *SSE1* and ER-UPR sensors during heat stress (Figure 2B, right panel). Upon treatment with Tm, *ire1*Δ or *hac1*Δ strains could not grow as expected, due to lack of mounting of ER-UPR (Figure 2C). Upon deletion of either *IRE1* or *HAC1*, the growth fitness observed for *sse1*Δ was completely abolished reiterating the fact that the Tm-resistance of *sse1*Δ strain depends on the presence of functional ER-UPR signalling by canonical Ire1-Hac1 pathway (Figure 2C). In sub-optimal concentrations of Tm that are not sufficient to mount ER stress, the phenotypes of *sse1*Δ or *ire1*Δ-*sse1*Δ and *hac1*Δ-*sse1*Δ remained similar to Tm-untreated condition (Figure 2D).

**Figure 2:**
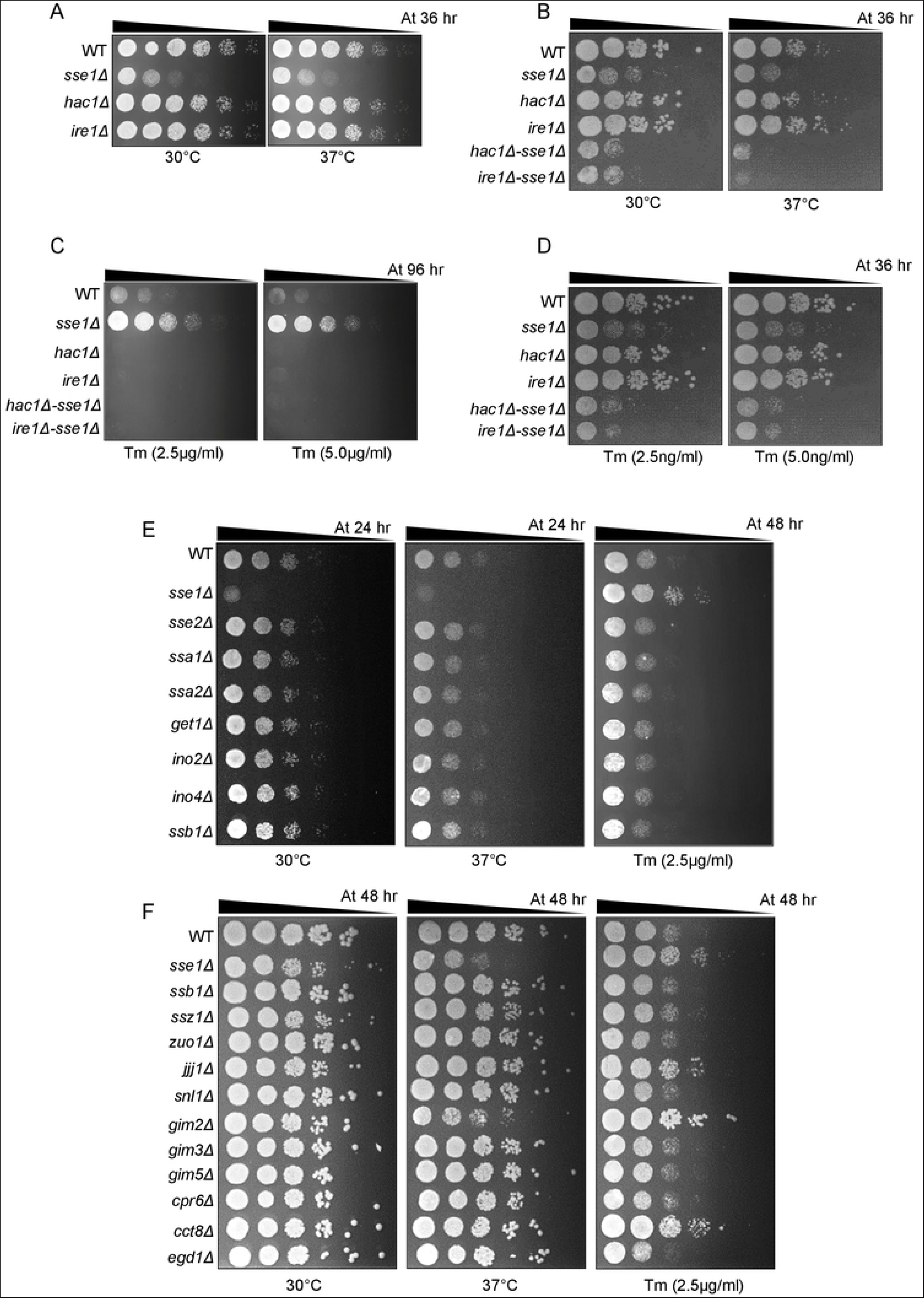
Tunicamycin resistance of *sse1Δ* is dependent on *IRE1-HAC1* pathway mediated ER-UPR signalling and the fitness is attributed to the abrogation of CLIPS function of Sse1 and not the basal high Heat Shock response of the strain. **A.** Yeast growth assay by serial drop dilutions using the strains WT, *sse1Δ*, *hac1Δ*, and *ire1Δ* in YPAD plates at permissive temperature (30°C) and at heat-shock condition (37°C). The time mentioned in hours represents the time of incubation before taking the image of the plates. **B**. Drop dilution assay as shown in panel A using the strains WT, and single deletion strains *sse1Δ*, *hac1Δ*, *ire1Δ*, and double deletion strains, *hac1Δ-sse1Δ* and *ire1Δ-sse1Δ* in YPAD plates at permissive temperature (30°C) and at heat-stressed condition (37°C). **C-D.** The same strains as in panel B were used for drop-dilution assay at optimal concentrations of Tm (2.5 µg/ml and 5.0 µg/ml) (panel **C**) and sub-optimal concentrations (2.5 ng/ml and 5.0 ng/ml) (panel **D**). **E.** Drop dilution assay of WT, *sse1Δ* and all other single deletion strains of yeast reported to exhibit high basal Heat Shock Response (HSR), (*sse2Δ*, *ssa1Δ*, *ssa2Δ*, *get1Δ*, *ino2Δ*, *ino4Δ*, and *ssb1Δ*) in YPAD plates at permissive temperature (30°C) (left panel), heat shock condition (37°C) (middle panel) and in presence of ER stressor Tm (2.5 µg/ml) (right panel). **F.** Similar to panel E, growth assay by serial drop dilutions was performed for the WT, *sse1Δ* and all other single deletion strains for CLIPS proteins apart from Sse1, (*ssb1Δ*, *ssz1Δ*, *zuo1Δ*, *jjj1Δ*, *snl1Δ*, *gim2Δ*, *gim3Δ*, *gim5Δ*, *cpr6Δ*, *cct8Δ*, and *egd1Δ*) in YPAD plates in permissive temperature (30°C) (left panel), heat shock condition (37°C) (middle panel) and in presence of an optimal concentration of tunicamycin (2.5 µg/ml) (right panel).

In summary, we show that *SSE1* genetically interacts with the ER-UPR pathway and Tm-resistance of *sse1*Δ strain is dependent on the efficient mounting of Ire1-Hac1 mediated ER-UPR signalling.

### Similar to *SSE1*, the deletion of three other CLIPS members, imparts resistance to tunicamycin-induced ER stress in yeast

It was previously shown that the deletion of many genes including the *sse1*Δ strain, increases the basal Heat Shock Response (HSR) of yeast, *S. cerevisiae* [15]. In another study, it was also known that Heat Shock Response (HSR) alleviates ER stress [16]. Thus, to explain the Tm-resistance of the *sse1*Δ strain, we hypothesized that the high basal heat shock response (HSR) of this strain during ER stress, may be responsible for conferring Tm-resistance.

To check this, we took the deletion strains of yeast from the YKO library which are reported to exhibit high basal HSR including *sse1*Δ. Among the eight deletion strains (*sse1*Δ, *sse2*Δ, *ssa1*Δ, *ssa2*Δ, *get1*Δ, *ino2*Δ, *ino4*Δ, and *ssb1*Δ) previously reported to show high basal HSR [15], none other than *sse1*Δ show any fitness during Tm-induced ER stress (Figure 2E). This data indicates that the growth fitness observed for the *sse1*Δ strain is not due to high basal HSR as none of the other deletion strains possessing high HSR exhibit any fitness during Tm-induced ER stress. Furthermore, to check the role of HSR, we expressed the constitutively active mutant of Hsf1 (Hsf1R206S) in WT and *sse1*Δ strains and checked the phenotype. Constitutive activation of HSR by Hsf1R206S in WT did not impart any fitness advantage to Tm-stress at 2.5µg/ml concentration of the stressor rather it increased the sensitivity to Tm-stress, negating the role of high HSR in Tm-resistance (Figure S2C, right panel). ln case of *sse1*Δ strain, Hsf1R206S expression leads to mild growth phenotype alleviation at ambient and at heat shock conditions, although there is no additional enhancement of tunicamycin resistance of the *sse1*Δ strain due to constant HSR activation (Figure S2C, right panel). This result further confirms Sse1’s specific role in cellular response during Tm-induced ER stress which cannot be complemented by activated HSR.

As Sse1 belongs to a class of chaperones termed as CLIPS (chaperones linked to protein synthesis) [10] due to its co-regulated expression pattern with cytosolic translation machinery, we hypothesized that deletion of *SSE1* would lead to altered or inefficient protein translation as indicated by enhanced sensitivity of *sse1*Δ strain to translation blocker, cycloheximide (Figure 1Bi and Bii). Thus, *SSE1* deletion may reduce the incoming protein load to ER which in consequence would help better management of ER stress, leading to the observed fitness. Next, we took the single deletion strains of all the CLIPS (*sse1*Δ, *ssb1*Δ, *ssz1*Δ, *zuo1*Δ, *jjj1*Δ, *snl1*Δ, *gim2*Δ, *gim3*Δ, *gim5*Δ, *cpr6*Δ, *cct8*Δ and *egd1*Δ) [10] and checked the phenotype during Tm-induced ER stress. Interestingly, apart from *sse1*Δ, we observed similar fitness in deletion strains of three other CLIPS namely *JJJ1*, *GIM2* and *CCT8* (Figure 2F). Jjj1 is a cytosolic J-domain co-chaperone protein of Hsp70 protein Ssa1 and it remains associated with large ribosomal subunits. Jjj1 was shown to be involved in the late stage, cytosolic steps of biogenesis of 60S ribosomal particles [17]. Gim2 is part of the prefoldin complex which works as co-chaperone of the CCT/TRiC chaperonin complex [18]. Prefoldin plays a crucial role in cytoskeleton assembly by helping the folding of actin and tubulin monomers [19]. Cct8 is part of the cytosolic chaperonin (barrel-shaped chaperone) CCT-ring complex [20]. So far, there is no evidence of interaction of *SSE1* with three of the CLIPS in literature, thus to check any genetic interaction between *SSE1* and these three CLIPS, we made double knockouts of *SSE1-JJJ1*, *SSE1-GIM2* and *SSE1-CCT8*. *SSE1* shows strong negative genetic interactions with both *JJJ1* and *GIM2* at a permissive temperature (30°C) (Figure S2D, left panel). *SSE1-GIM2* genetic interaction is very strong at permissive temperatures and the double deletion is synthetically lethal during heat stress at 37°C (Figure S2D, middle panel). Interestingly, in the double deletion strains of *SSE1-JJJ1* and *SSE1-GIM2,* Tm-resistance of individual single deletion strains of these CLIPS is abolished. The double knock-out strains show similar Tm-sensitivity like wild type yeast (Figure S2D, right panel). This data indicates that during Tm-induced ER stress, *SSE1* genetically interacts with *JJJ1* and *GIM2* individually and the Tm-resistance of *sse1*Δ depends on intact function of *JJJ1* or *GIM2* and vice versa. *SSE1-CCT8* double knockout strain shows similar fitness like the single deletion indicating no genetic interaction of *SSE1* and *CCT8* during tunicamycin induced ER stress (Figure S2D).

In summary, we found that the absence of CLIPS function of Sse1 rather than the high heat shock response of *sse1*Δ strain plausibly imparts resistance to Tm-induced ER stress.

### Sse1 controls the ER-stress-induced changes in cellular protein translation

In the previous section, we have shown that in absence of *SSE1* or individual deletion of the other three CLIPS, *JJJ1*, *GIM2* and *CCT8*, imparts tunicamycin resistance to yeast cells. Thus, it was interesting to assess any changes in protein translation following Tm-induced ER stress, especially in the absence of Sse1. To capture the status of cellular translating ribosomes, we performed the polysome profiling of WT and *sse1*Δ strain in the absence of any external stress and following imparting Tm-induced ER stress. In the physiological condition, the polysome profile of WT yeast cells shows a large part of total ribosomes in the polysome fraction along with the monosome (80S) fraction (Figure 3A, left panel). Upon inducing ER-stress by tunicamycin, a significant part of the polysomes is shifted to monosome fraction in WT yeast cells (Figure 3A, middle panel). In case of *sse1*Δ strain, the polysome profile reveals a similar presence of polysome and monosome fractions (Figure 3A, left panel). Interestingly, upon Tm-treatment, an equivalent reduction in polysome fraction is not explicitly prominent in the *sse1*Δ strain in contrast to WT cells, although the peak of monosome is prominently enhanced indicating a possibility of more ribosome assembly as monosomes in the *sse1*Δ strain during Tm-induced stress (Figure 3A, right panel). This data indicates an important regulatory role of Sse1 during ER-stress-induced re-organization of the cellular translation apparatus. This finding points towards the possibility of unhindered polysome and monosome-driven protein translation in *sse1*Δ strain compared to WT cells during ER stress.

**Figure 3:**
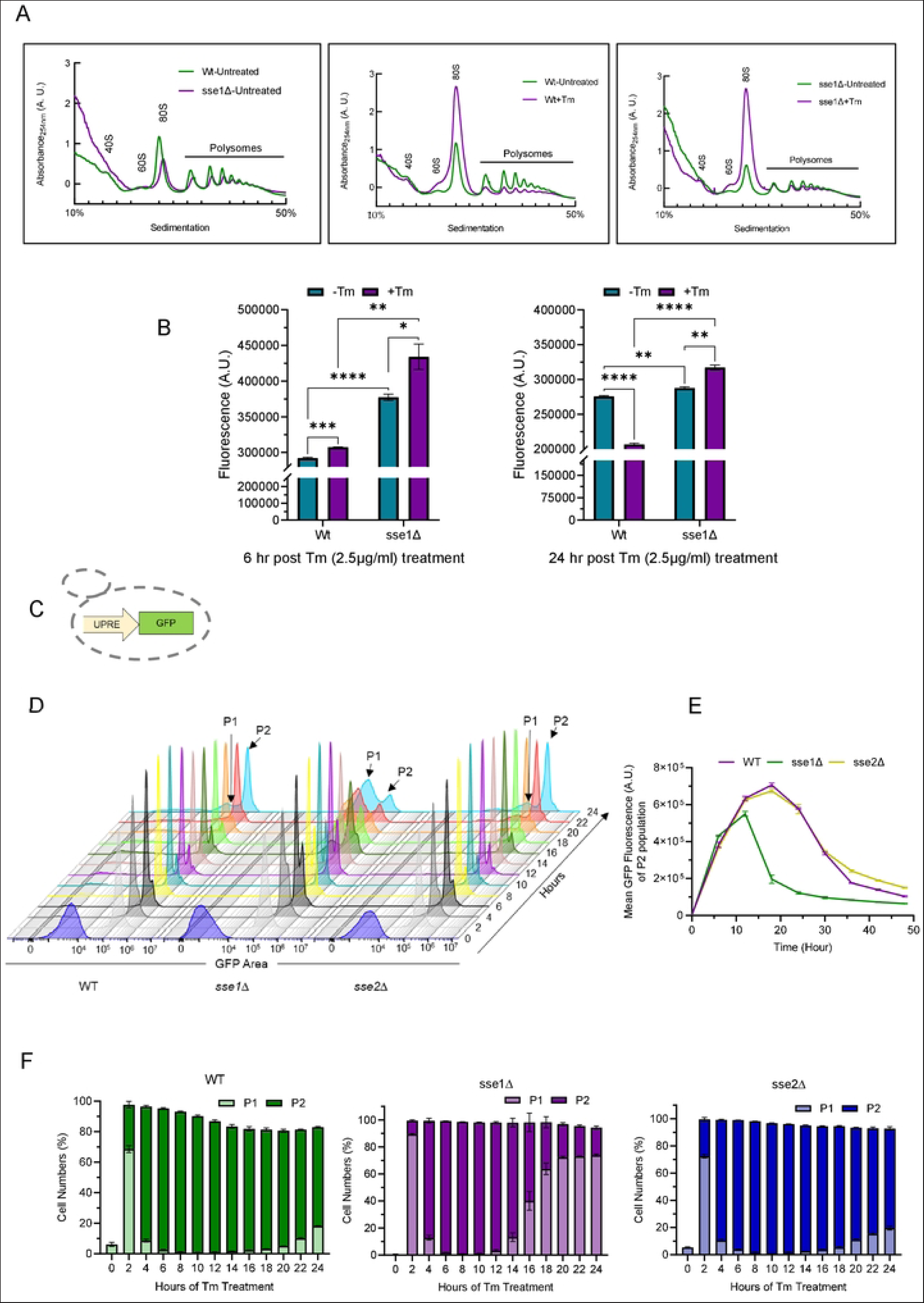
Sse1 plays an important role in modulating tunicamycin-induced ER stress-associated changes in protein translation. **A.** Polysome profiles are plotted for the WT and *sse1Δ* strains isolated from untreated cells and cells treated with Tm (2.5 µg/ml) and comparison between polysome profiles of (left panel) untreated WT vs untreated *sse1Δ*, (middle panel) untreated vs Tm-treated WT cells and (right panel) untreated vs Tm-treated *sse1Δ* cells are shown. **B.** The rate of translation of WT, and *sse1Δ* strains in untreated and Tm (2.5 µg/ml) treated conditions were analyzed using the CLICK-IT chemistry reaction using L-Azido-homoalanine and Alexa-Fluor 488 alkyne dye. The incorporated fluorescence in newly synthesized proteins was measured in each sample by flow cytometry and was plotted as a bar plot with whisks representing SEM. The left panel shows the incorporated fluorescence following 6 hours of Tm-stress in comparison to untreated cells of WT and *sse1Δ* strains. The right panel shows same after 24 hours of Tm-treatment. Statistical significance was calculated using unpaired T-tests and the pairwise comparison outputs were plotted in the graph (Left Panel: WT-Tm/WT+Tm, two-tailed p=0.0003, ***; *sse1*Δ-Tm/*sse1*Δ+Tm, two-tailed p=0.0367, *; WT-Tm/*sse1*Δ-Tm, two-tailed p<0.0001, ****; WT+Tm/*sse1*Δ+Tm, two-tailed p=0.0021, **. Right Panel: WT-Tm/WT+Tm, two-tailed p<0.0001, ****; *sse1*Δ-Tm/*sse1*Δ+Tm, two-tailed p=0.0019, **; WT-Tm/*sse1*Δ-Tm, two-tailed p=0.0052, **; WT+Tm/*sse1*Δ+Tm, two-tailed p<0.0001, ****). **C.** Schematic of the yeast strain YMJ003 (wild type) that serves as a reporter strain for ER-UPR activation. The strain contains the GFP under Unfolded Protein Response Element (UPRE) to report for ER-UPR induction. **D.** Kinetics of ER-UPR activation was measured by measuring UPRE-GFP fluorescence by flow cytometry and shown as combined overlaid histograms where each histogram represents the data of the strains WT (YMJ003), *sse1Δ*, and *sse2Δ* (in YMJ003 strain background) at designated time points following treatment with Tm (2.5 µg/ml). The shift in the two populations P1 (basal state) and P2 (UPR-activated population), among the strains are marked accordingly. **E.** The P2 population’s (marked in the previous panel) mean GFP fluorescence intensity over time is plotted in this line plot for the same set of WT, *sse1*Δ and *sse2*Δ cells. The whisks over each time point value represents SEM. **F.** The kinetic change in the number of cells of the two population, P1 & P2, marked in the previous panel D are represented as stacked columns over time for the same set of WT, *sse1*Δ and *sse2*Δ cells. The whisks over each time point value represents SEM.

To check this, we measured the synthesis of new proteins by incorporating AHA (L-Azidohomoalanine) for tagging the newly synthesized proteins by Click-IT chemistry [21]. We measured the new protein synthesis status at multiple time points following the induction of ER stress by Tm-treatment using flow cytometry. Interestingly, after 6 hours of Tm-treatment, the amount of newly translated proteins in *sse1*Δ strain was significantly higher than the WT cells which nicely corroborated with the ribosome profile of these two strains (Figure 3B, left panel). When we measured the AHA fluorescence post 24 hours of Tm-treatment, compared to untreated cells, we observed a significant reduction in the incorporated fluorescence in WT cells indicative of a reduction of new protein synthesis (Figure 3B, right panel). In sharp contrast, Tm-treated *sse1*Δ cells showed significantly higher new protein synthesis compared to untreated cells at the same time points (Figure 3B, left and right panels). Finally, the comparison of new protein synthesis status between WT and *sse1*Δ strains post 24 hours of Tm-stress showed a drastic increase in AHA incorporation in *sse1*Δ cells indicative of significantly higher translation of new proteins in *sse1*Δ strain. This trend of new protein synthesis remained similar upon continuing the Tm-stress for 48 hours (Figure S3A).

As AHA incorporation indicates all new protein translation, to exclusively measure the formation of UPR-induced proteins following ER stress, we employed the previously described UPR-reporter strain, YMJ003 [22, 23]. This strain contains genome-integrated EGFP (<u>E</u>nhanced <u>G</u>reen <u>F</u>luorescence <u>P</u>rotein) which is expressed under the UPR element (UPRE) whenever cells experience ER stress (Figure 3C) [22, 23]. In the background of YMJ003, we deleted *SSE1* and also *SSE2* (as control) which showed similar phenotypes as observed in the case of BY4741 WT background as described in both Figure 1 and 2 (resistance against optimal concentrations of Tm sufficient to elicit ER-UPR and absence of growth fitness in sub-optimal Tm concentrations) (Figure S2A and S2B). When we compared the kinetics of UPR by following the GFP fluorescence by flow cytometry, we observed a very distinctive kinetics of ER-UPR induction in *sse1*Δ strain as compared to WT or *sse2*Δ strains (Figure 3D and 3E). In all cases, we observed the time-dependent appearance of a high-fluorescent population (UPR-activated population denoted as P2 population) after the Tm-stress. After 2 hours of Tm-treatment, about 20-30% population shifts to the P2 population which becomes nearly 100% by 6 hours in all the three strains indicating near complete mounting of ER-UPR (Figure 3F). Interestingly, for the *sse1*Δ strain, we observed a faster reversal to basal state (P1 population) compared to WT or *sse2*Δ strains (Figure 3D and 3F, middle panel). By 16 hours post-Tm-treatment, we observed around 50% of *sse1*Δ cells in the P1 population whereas WT or *sse2*Δ strains showed less than 5% cells in the P1 population (Figure 3D, 3F and Figure S3B). By 24 hours, almost 75% of *sse1*Δ cells shift to the P1 population while only about 18% of the WT or *sse2*Δ cells are present in the P1 population (Figure 3D, 3F and Figure S3B). This result nicely indicates a faster reversal from the UPR-activated state for *sse1*Δ strain in comparison to WT strain. The mean GFP intensity plot of the UPR-activated cell population (P2 population) of these 3 strains further indicates a quicker response to ER stress and reversal to the basal state by *sse1*Δ strain as compared to WT or *sse2*Δ strains (Figure 3E). WT or *sse2*Δ strains show much higher and sustained response to chronic Tm-induced ER stress (Figure 3E). This result shows the important role of Sse1 in maintaining a prolonged response to ER stress during global ER stress by tunicamycin.

Next, to follow the actual cellular response in terms of synthesis of UPR-responsive proteins following induction of ER stress, we took different GFP-tagged strains of ER-UPR target proteins like Pdi1, Lhs1, Sec62 and Ubc7 to monitor the protein synthesis following induction of ER stress, by flow cytometry (Figure 4A, upper panel). As a control of non-ER-UPR target protein, we took the Tdh1-GFP strain. In these GFP-tagged strains, we deleted the genomic copy of *SSE1* (Figure 4A, lower panel). Interestingly, for all ER-UPR targets Pdi1, Lhs1, Sec62 or Ubc7, we observed significant time-dependent increased expression of these proteins following Tm-treatment indicating mounting of ER-UPR followed by a gradual decrease in the protein level indicative of decay of the protein levels and the ER-stress response (Figure 4B-E). The expression of these proteins remained at a much lower basal level in the untreated cells confirming the suitability of checking the expression levels of these target proteins as reporters of ER-UPR induction by tunicamycin (Figure 4B-E). Importantly, the corresponding *SSE1*-deleted versions of the GFP-tagged strains showed the highest expression of all these ER-UPR target proteins around 18-22 hrs post-Tm-treatment followed by a decrease in the protein expression to basal level by 30-32 hrs (Figure 4Bi-Ei). In contrast, in case of the WT strains, the expression of these proteins peaked at later time points (28-30 hrs post-Tm-treatment) followed by a decay in the protein levels at much later time points (around 40 hrs) (Figure 4Bi-Ei). Similar to the UPRE-GFP reporter, the GFP-tagged ER-UPR target proteins also showed a UPR-activated state (P2 population) at higher fluorescence intensity and a lower fluorescent P1 population in the later time points indicative of basal state after reversal from the UPR-activated state (Figure 4Bii-Eii). Overall, the quick response to ER stress and faster reversal to the basal state from the UPR-activated state by the *sse1*Δ strain in comparison to the WT strain as detected by the UPRE-GFP reporter described in the previous section (Figure 3D-F) was further confirmed by following the UPR-activated expression of actual cellular UPR-target proteins (Figure 4B-E). The control protein Tdh1 did not show any ER-stress mediated increased expression as expected (Figure 4Fi and Fii), further confirming that the time-dependent overexpression of the UPR-target proteins Pdi1, Lhs1, Sec62 and Ubc7 as described above, was exclusively due to induction of ER-UPR by tunicamycin.

**Figure 4:**
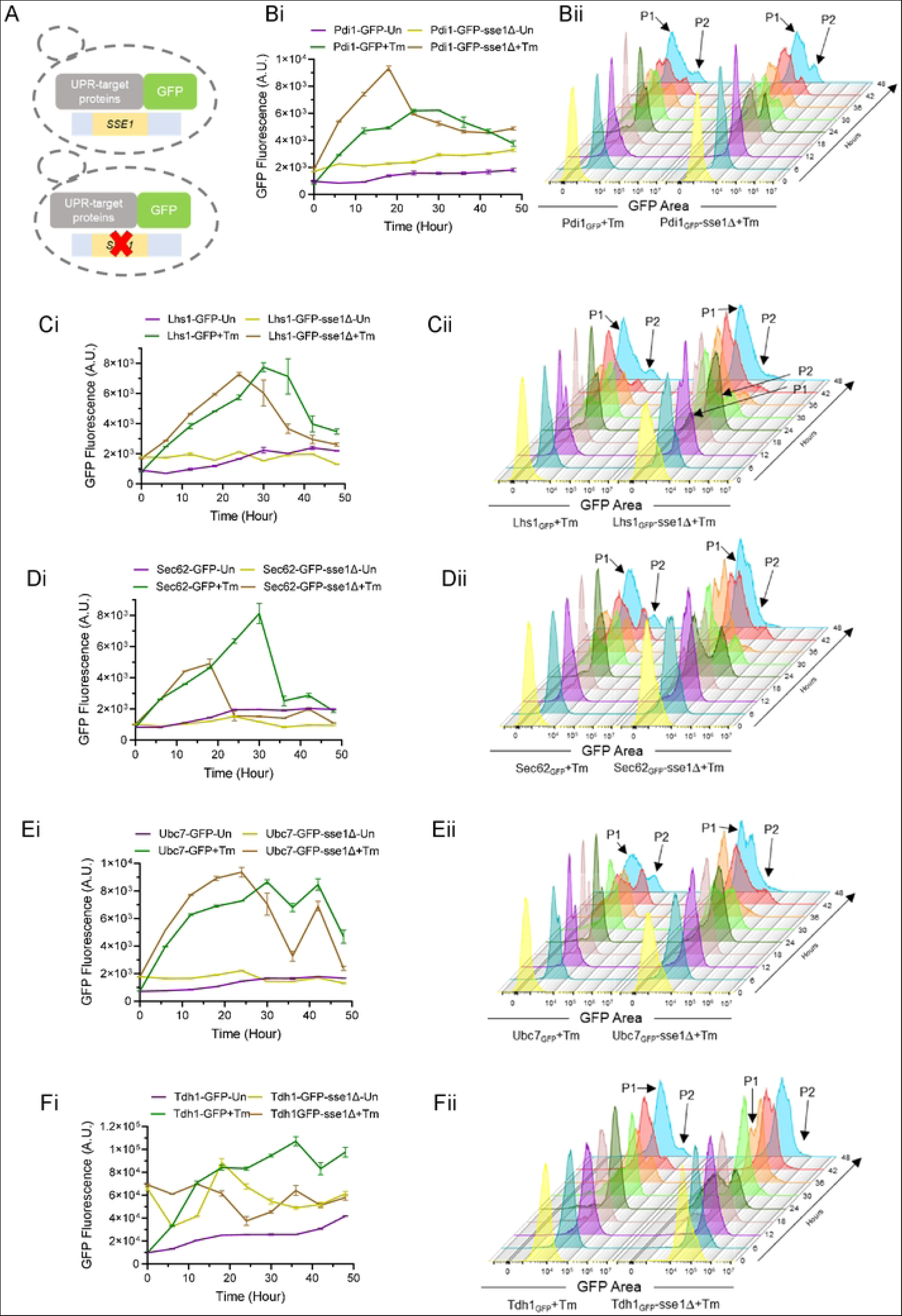
The ER-UPR kinetics is different in absence of Sse1. **A.** (left) Schematic of the yeast strains where individual UPR target genes are tagged with GFP protein, so that their real time translation can be monitored by measuring GFP fluorescence. (right) Using these strains as background, *SSE1* was deleted using *URA3* cassette in each of the GFP reporter strain. The left panel strains serve as the WT cells and the right panel strain serve as *sse1*Δ strain. **B-F.** The GFP tagged UPR target gene’s expression over time are plotted as line graphs **(Bi-Fi)** along with combined overlaid histograms **(Bii-Fii)** where each histogram represents the data of the respective strains at designated time points following treatment with with Tm (2.5 µg/ml). The line plots of the expression of cellular target genes of ER-UPR like Pdi1-GFP **(Bi)**, Lhs1-GFP **(Ci)**, Sec62-GFP **(Di)** and Ubc7-GFP **(Ei)** strains with *sse1*Δ counterparts along with untreated and Tm-treated samples are shown. The combined overlaid histograms of the same above-described ER-UPR targets that are Pdi1-GFP **(Bii)**, Lhs1-GFP **(Cii)**, Sec62-GFP **(Dii)** and Ubc7-GFP **(Eii)** strains with *sse1*Δ counterparts which are Tm-treated samples are shown. **F. (Fi)** Line plots of the Tdh1-GFP strain (non-ER-UPR control) with *sse1*Δ counterparts along with untreated and Tm-treated samples are shown. **(Fii)** Combined overlaid histograms of the Tdh1-GFP strain (non-ER-UPR control) with *sse1*Δ counterparts that are Tm-treated are shown here.

In summary, we show that in WT cells, in the presence of Sse1, the unfolded protein response is a sustained process following Tm-induced ER stress. Interestingly, in absence of Sse1, activation of ER-UPR as well as reversal to basal state indicative of restoration of homeostasis following Tm-induced ER stress, is much quicker which can explain the fitness to Tm-stress observed in the *sse1*Δ strain.

### Cellular response to Tm-induced ER stress is distinctly different in the absence of Sse1

To understand the cellular response during Tm-stress, we did an RNA sequencing-based transcriptome analysis and a label-free quantitative proteomics analysis of untreated and Tm-treated WT and *sse1*Δ strains. For transcriptome analysis, along with 40 other yeast samples of similar genetic background, WT and *sse1*Δ strains were subjected to RNA sequencing in untreated and Tm-treated conditions. To identify the genes that are differentially upregulated or downregulated at a particular sample, the Z-score of expression for each gene across all these yeast strains was calculated as described previously [24]. The genes above or below Z-score 2 at a particular condition were considered as significantly upregulated or downregulated, respectively. Using Z-score analysis of the transcriptomics data, we found 885 genes to be differentially upregulated and 9 genes to be downregulated in the WT strain upon Tm-treatment. In contrast, *sse1*Δ strain showed 415 genes to be upregulated and 115 genes to be downregulated in the untreated condition. Upon Tm-treatment, *sse1*Δ strain showed 370 genes to be upregulated and 14 genes to be downregulated. Next, we did a pathway enrichment analysis of upregulated genes (Figure 5) found in the WT-treated cells and *sse1*Δ cells in both untreated and Tm-treated conditions. Among the top 30 enriched pathways in Tm-treated WT cells, response to unfolded proteins, macromolecule glycosylation, protein glycosylation, ERAD pathway etc. clearly shows the response to tunicamycin stress and activation of ER-UPR (Figure 5A and Table S1). Many enriched pathways show changes in intracellular protein trafficking and protein localization. In the *sse1*Δ strains, the untreated condition shows a very significant enrichment of ribosome assembly, cytoplasmic protein translation and various biosynthetic pathways (Figure 5B and Table S1). In the *sse1*Δ Tm-treated transcriptome, we found macromolecule and protein glycosylation, and protein quality control pathways similar to WT-treated cells (Figure 5C and Table S1). Interestingly, we found cytoplasmic translation, cytoskeleton reorganization and cell cycle pathways to be uniquely enriched in the *sse1*Δ treated cells (Figure 5C and Table S1).

**Figure 5:**
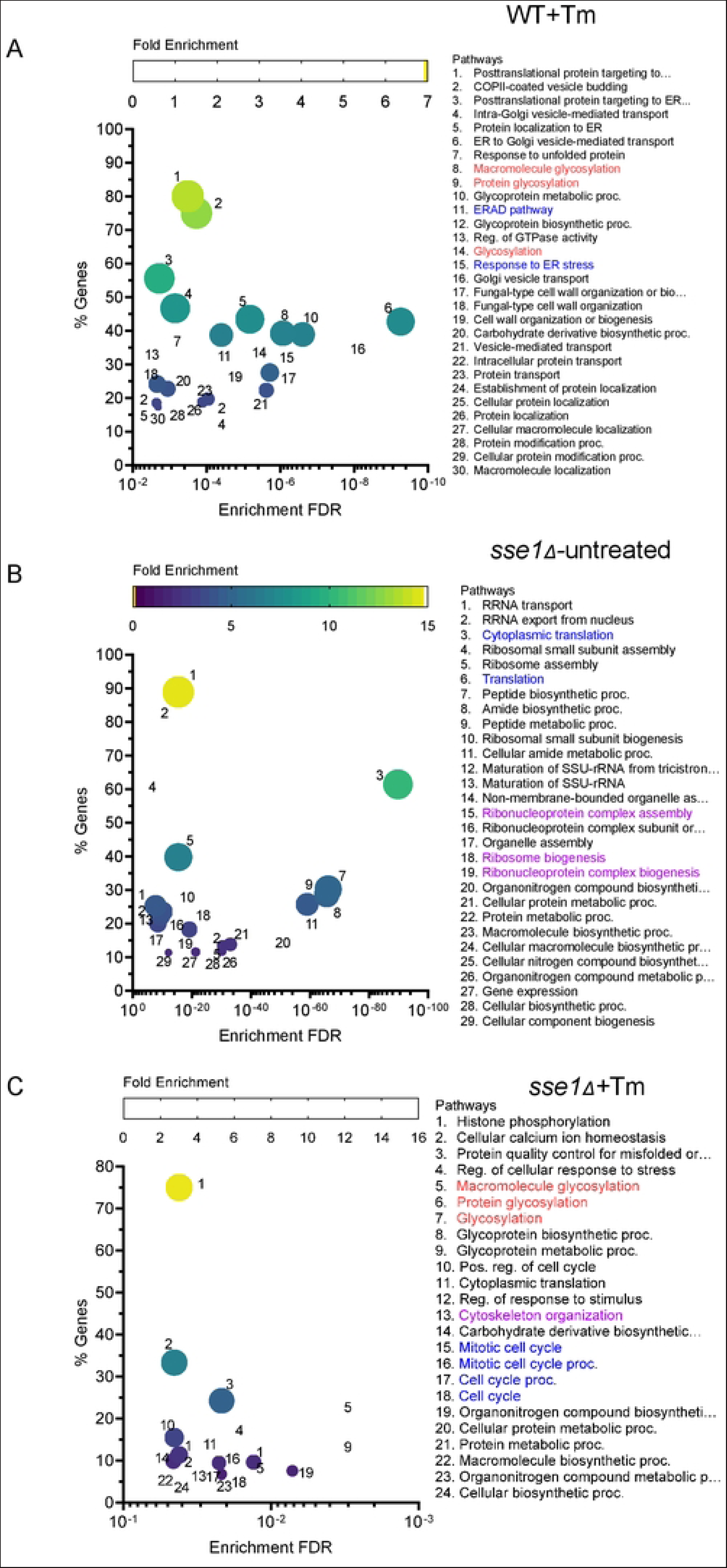
The pathway enrichment analysis of the cellular transcriptome upon tunicamycin stress in WT and *sse1*Δ strains. **A-C.** Transcriptome analysis of WT and *sse1*Δ strains in untreated condition and after treatment with tunicamycin (2.5 μg/ml) along with other yeast strains of same genetic background as described previously [24] was done by RNA sequencing. The outputs were converted to Z scores. Transcripts above and below z-score 2 were considered as differentially upregulated or downregulated, respectively. The multivariate bubble plots showing the enriched upregulated pathways with the attributes - the Enrichment False Discovery Rate (Enrichment FDR) on the X-axis, the percentage of genes identified for each of the enriched pathway with respect to the total annotated genes on the Y-axis, the colour scale represents the Fold Enrichment, and the larger bubble size signifies highly enriched and significant pathways. **A.** Bubble plot representing the pathways enriched in WT cells when treated with tunicamycin (2.5 μg/ml). Bubble plot showing the enriched pathways of the *sse1*Δ strain in untreated condition (panel B) and tunicamycin (2.5 μg/ml)-treated condition (panel C). The important enriched pathways in all three cases have been highlighted in different colors.

Next, by quantitative proteomics analysis, 904 proteins were detected in all three replicates of WT and *sse1*Δ Tm-treated and untreated samples which were analyzed further. Comparison of protein levels of untreated *sse1*Δ cells with respect to WT cells at basal condition (no stress) revealed 12 and 23 proteins to be differentially upregulated and downregulated, respectively (Figure 6A and Table S2). Rest 869 proteins did not show any significant changes in the expression level. Among the upregulated proteins in untreated *sse1*Δ cells with respect to WT cells, another cytosolic NEF, Fes1, was detected indicating cellular response to compensate for the loss of Sse1’s NEF activity (Figure 6B). Comparison of protein levels of Tm-treated WT cells with respect to untreated WT cells revealed a drastic increase in the number of differentially expressed proteins due to ER stress (Figure 6C, Figure S3C and Table S2). Tm-induced stress led to differential upregulation of 49 proteins and downregulation of 88 proteins in the WT strain and changes in expression of the rest of the 767 proteins remained insignificant (Figure 6A). Among the upregulated proteins, ER-UPR responsive proteins like Kar2 and Pdi1 were found along with many other differentially overexpressed proteins involved in the maintenance of proteostasis like Trx3, Grx2, Hsp42 etc. (Figure 6C and Figure S3C). Interestingly, when we compared the protein levels of Tm-treated *sse1*Δ cells with untreated *sse1*Δ cells, the number of upregulated proteins was considerably higher (105 proteins) and the number of downregulated proteins (50 proteins) was much less compared to WT cells under the same condition (Figure 6A). Like WT cells, *sse1*Δ cells also showed differentially upregulated Kar2, Pdi1, Trx3, and Grx2 upon Tm-treatment (Figure 6D and Figure S3C). The more number of overexpressed proteins in *sse1*Δ cells during Tm-stress reiterated that protein translation is less inhibited in *sse1*Δ in comparison to WT cells upon Tm-induced ER stress as shown by AHA-incorporation by Click-IT reaction (Figure 3B, left panel). Interestingly, a comparison of Tm-treated *sse1*Δ cells with respect to Tm-treated WT cells showed the upregulation of 48 proteins and the downregulation of 11 proteins. Among the upregulated proteins, both subunits of yeast NAC (Nascent polypeptide Chain Associated Complex), Egd1 (yeast orthologue of NACβ) and Egd2 (yeast orthologue of NACα) [25] were present (Figure 6E). The differential overexpression of NAC subunits in *sse1*Δ cells during ER stress indicates a cellular response to protect the newly synthesized proteins as NAC protects the nascent chains (NC) from aggregation and misfolding upon emerging from ribosome exit tunnels [25, 26]. This data is in corroboration to unchanged level of ribosome-bound-chaperone, Ssb1, a wellknown chaperone that binds the NCs upon emergence from ribosome-exit tunnels, in *sse1*Δ cells following Tm-induced ER stress (Figure S3D). In contrast, WT cells show a prominent reduction of Ssb1-bound to ribosomes following Tm-treatment (Figure S3D). Thus, it is distinctly evident from multiple results that *sse1*Δ strain more efficiently continues protein translation during Tm-induced ER stress.

**Figure 6:**
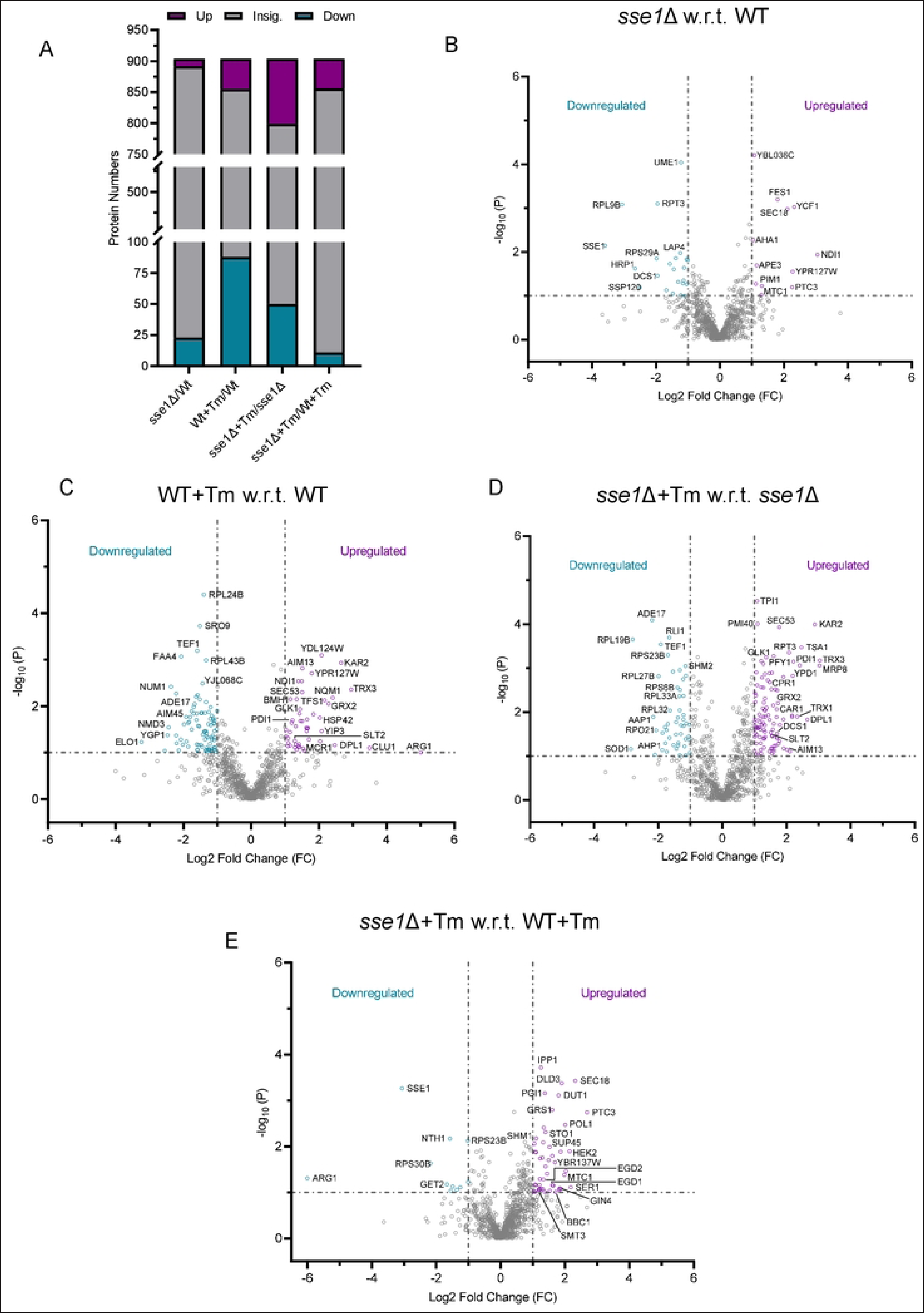
The changes in cellular proteome upon tunicamycin stress in WT and *sse1Δ* strains. **A.** The total number of proteins identified by quantitative mass spectrometry and the outputs of the statistical analysis of various pair-wise comparisons are plotted as composite bar plots showing the total number of proteins that were differentially upregulated, downregulated and insignificant for each of the comparisons between the *sse1Δ* and WT strain in untreated and after treatment with optimal tunicamycin (2.5 µg/ml). **B-E.** Volcano plots showing the differentially expressed (upregulated and downregulated) as well as proteins with insignificant changes in the expression level in the case of untreated *sse1Δ* strain with respect to the untreated WT strain (panel **B**), in the case of Tm (2.5 µg/ml) treated WT strain with respect to the untreated WT strain (panel **C**), in the case of Tm (2.5 µg/ml) treated *sse1Δ* strain with respect to the untreated *sse1Δ* strain (panel **D**) and in case of Tm (2.5 µg/ml) treated *sse1Δ* strain with respect to Tm (2.5 µg/ml) treated WT strain (panel **E**).

### Sse1 plays a crucial role in controlling ER-stress-induced cell division arrest and cell viability

As *sse1*Δ strain showed prominent growth fitness during long-standing Tm-stress and tunicamycin stress is known to cause cell division arrest in yeast [27, 28], it was interesting to check the status of cell division of this deletion strain during Tm-induced ER stress. Furthermore, we found that mitotic cell cycle and cell cycle pathways to be significantly upregulated in Tm-treated *sse1*Δ cells by pathway enrichment analysis of the transcriptome data, as described before (Figure 5C). Thus, it prompted us to check the cellular morphology and cell cycle status of the Tm-untreated and treated WT and *sse1*Δ cells. To check the cellular morphology, we performed imaging of yeast cells by confocal microscopy at different time points following Tm-stress. WT cells showed significantly higher cell size (3 to 4 times) and granularity in the later time points of stress (most prominent from 18 hrs post-Tm-treatment) compared to *sse1*Δ cells (Figure 7A and 7B). In the *sse1*Δ strain, the increase in cell size post-Tm-stress was also observed but to a much lesser extent compared to WT cells (Figure 7A and 7B). This data indicated that there is a block in cell division in WT cells as a response to ER stress which is bypassed in the *sse1*Δ strain. To validate this finding, we did a cell cycle analysis with Sytox Green dye as described before [29]. The untreated cells in both WT and *sse1*Δ strains showed cells possessing 1C and 2C DNA content. After 6 hrs of Tm-treatment, cell cycle analysis showed the appearance of populations with more DNA content (3C and 4C) in both the strains (Figure 7C, 7D and Figure S4Ai and S4Aii) indicating cytokinesis arrest. Upon long-standing stress for 24 hrs of Tm-treatment, all the cells of the WT strain were populated in a higher DNA-containing population (3C, 4C) indicating a major block in cell division (Figure 7E and Figure S4Aiii). In sharp contrast, after 24 hours of Tm-stress, *sse1*Δ strain showed a majority of the cells in the 1C and 2C population almost overlapping with the untreated condition (Figure 7F and Figure S4Aiv) indicating progression of cell division. This data is intriguing and indicates an important role of Sse1 in controlling cell division arrest during Tm-induced ER stress.

**Figure 7:**
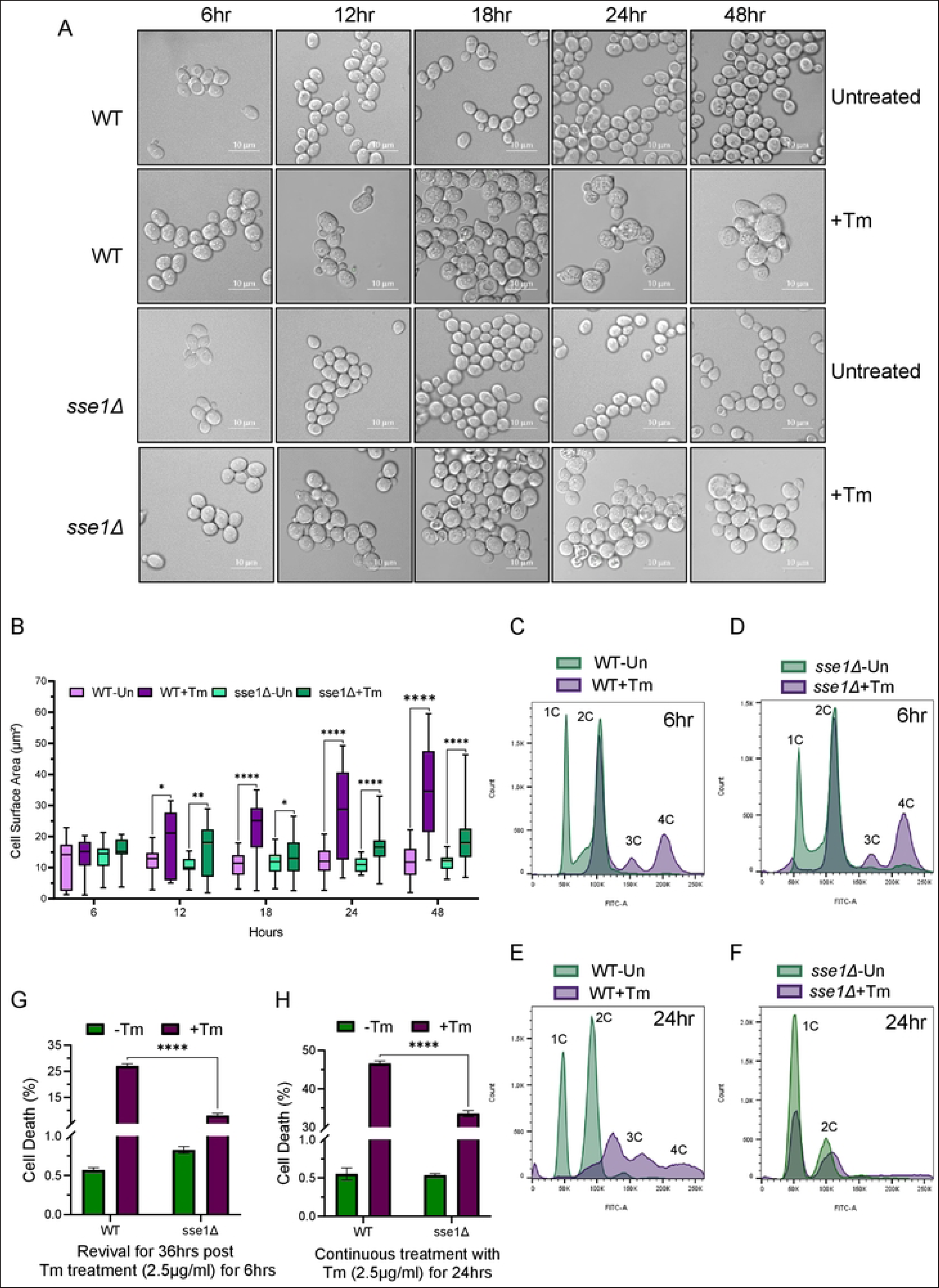
Sse1 controls tunicamycin-induced ER stress mediated cell division arrest and cell death of yeast. **A.** Images taken by confocal microscopy of WT and *sse1Δ* cells following treatment with Tm (2.5 µg/ml) along with the untreated control cells. Images were captured at 6-, 12-, 18-, 24-, and 48-hours’ time point respectively. **B.** The quantitation of the size (surface area in μm^2^) of the yeast cells from the previous panels which are represented as a box and whisks plot where the top whisk representing highest and the bottom whisk representing lowest individual cell surface area. The horizontal line within each box represents the median cell surface area. Statistical significance was calculated using unpaired T-tests and the pairwise comparison outputs were plotted in the graph (12 Hours: WT-Tm/WT+Tm, two-tailed p=0.0144, *; *sse1*Δ-Tm/*sse1*Δ+Tm, two-tailed p=0.0060, **; 18 Hours: WT-Tm/WT+Tm, two-tailed p<0.0001, ****; *sse1*Δ-Tm/*sse1*Δ+Tm, two-tailed p=0.0403, *; 24 Hours: WT-Tm/WT+Tm, two-tailed p<0.0001, ****; *sse1*Δ-Tm/*sse1*Δ+Tm, two-tailed p<0.0001, ****; 48 Hours: WT-Tm/WT+Tm, two-tailed p<0.0001, ****; *sse1*Δ-Tm/*sse1*Δ+Tm, two-tailed p<0.0001, ****). **C-F.** Cell cycle analysis was done for the WT and *sse1Δ* cells using the DNA binding fluorescent dye Sytox Green following treatment with Tm (2.5 µg/ml) with untreated controls. The data were captured after 6 hours and 24 hours of Tm (2.5 µg/ml) treatment. **C.** The overlaid histogram represents the pairwise comparison of WT-untreated/WT+Tm cell cycle pattern at 6hours post Tm treatment. **D.** The overlaid histogram represents the pairwise comparison of *sse1Δ*-untreated/*sse1Δ*+Tm cell cycle pattern at 6 hours post Tm treatment. **E.** The overlaid histogram represents the pairwise comparison of WT-untreated/WT+Tm cell cycle pattern at 24 hours post Tm treatment. **F.** The overlaid histogram represents the pairwise comparison of *sse1Δ*-untreated/*sse1Δ*+Tm cell cycle pattern at 24 hours post Tm treatment. **G.** Cell death percentage was analysed using propidium iodide (PI) staining through flow cytometry and was plotted as a bar plot (with whisks representing SEM, n=3) using the strains WT (BY4741) and *sse1*Δ (in BY4741 strain background) at the revival stage [after 36 hours following optimal Tm (2.5 µg/ml) treatment for 6 hours]. Statistical significance was calculated using unpaired T-tests and the significant pair was plotted in the graph (WT+Tm/*sse1*Δ+Tm, p<0.0001, ****). **H.** A similar cell death percentage as shown in panel G was determined for WT (BY4741) and *sse1*Δ (in BY4741 strain background) after chronic ER stress of 24 hours by Tm (2.5 µg/ml) treatment. Statistical significance was calculated using unpaired T-tests and the significant pair was plotted in the graph (WT+Tm/*sse1*Δ+Tm, p<0.0001, ****).

To test whether the escape of cell division arrest of *sse1*Δ strain during Tm-stress leads to alteration in cell viability following acute or chronic ER stress, we measured the cell viability using Propidium Iodide (PI) staining of yeast cells. In the case of short-term Tm stress for 6 hours followed by recovery for 36 hours, it showed ~27% cell death in WT cells (Figure 7G). In contrast, cell death was significantly less (~8%) in the case of *sse1*Δ cells strain (Figure 7G). In case of uninterrupted chronic Tm-treatment, the percentage of cell death of *sse1*Δ cells was always significantly lower compared to WT cells (Figure 7H and S4B). These data together indicate that the Tm-resistance of the *sse1*Δ strain observed in growth assays is due to evasion of cell division arrest and significantly higher cell survival of the *sse1*Δ cells during Tm-induced ER stress. The role of Sse1 in controlling the cell division remains elusive at the moment. Altogether, we show an important role of Sse1 in preventing cell division and successful triggering of cell death pathways following Tm-induced ER stress.

## Discussion

The Hsp110 group of molecular chaperones are exclusively found in eukaryotes although it’s Hsp70 partners are conserved across almost all kingdoms of life. As a representative member of Hsp110s, the cellular roles of yeast Hsp110, Sse1, have been extensively explored for more than the past two decades. Although Sse1’s role as a potent NEF of cytosolic Hsp70s (Ssa and Ssb) is well documented, it’s individual role in protein homeostasis beyond co-chaperone activity, if any, is not much explored. In this work, we reveal a yet unexplored role of Sse1 in regulating ER-unfolded protein response during ER stress. We show that Sse1 is required for an optimum cellular response during overwhelming ER stress caused by tunicamycin (Tm), an N-linked glycosylation inhibitor. Our data reveal that in the absence of Sse1, cells acquire an unusual resistance to ER stress. Importantly, this Tm-resistance of *sse1Δ* strain is heavily dependent on the successful induction of Ire1-Hac1 signalling mediated ER-UPR. To explain the Tm-resistance, we initially hypothesized a multitude of possibilities like; 1) high basal heat shock response (HSR) of *sse1Δ* strain, 2) decreased load of newly synthesized ER proteins due to the absence of CLIPS function of Sse1 which could be beneficial during ER stress. In contrast, recent literature has shown that the efficiency of ER-reflux of proteins is reduced in the absence of Sse1 during Tm-stress [11], which may increase the load of aberrantly folded proteins inside ER and compromise the ER protein homeostasis leading to higher basal ER-UPR in *sse1*Δ cells. In that scenario the cellular fitness of yeast in the absence of Sse1 is counter-intuitive.

To find out the most probable explanation of the Tm-resistance of the *sse1Δ* strain, we explored further and ruled out the contribution of high basal HSR as other strains possessing high basal HSR or constitutive activation of HSR do not show similar Tm-resistance. Interestingly, deletion strains of three other CLIPS, *JJJ1*, *GIM2* and *CCT8* apart from *SSE1* show the same Tm-resistance phenotype indicating the importance of the absence of individual CLIPS function as one of the common factors to gain Tm-resistance. This result prompted us to explore the role of Sse1 in modulating the protein translation status during Tm-induced ER stress. We found that Sse1 plays a crucial role in the stress-induced reorganization of the majority of translating ribosomes from polysomes to monosomes. In the absence of Sse1, this ribosomal reorganization is inefficient leading to a less prominent reduction in polysome fraction and an additional gain in the monosome fraction by a yet unknown mechanism. Together such ribosomal status in the *sse1*Δ cells leads to continued protein translation during longstanding Tm-induced ER stress, in complete contrast to ceased protein translation in WT cells. Our initial hypothesis was that the continuation of protein translation leads to a more efficient synthesis of ER-UPR-induced genes leading to better ER-stress management by *sse1Δ* strain. To check this, when we monitored the ER-UPR kinetics, we found the kinetics of UPR is prominently different from WT cells. The *sse1Δ* strain shows faster activation as well as faster reversal from the UPR-activated state. This result indeed shows that *sse1Δ* strain can activate the ER-UPR faster and restore the homeostasis quicker than WT leading to the fitness advantage during Tm-stress (summarized schematically in Figure 8).

**Figure 8:**
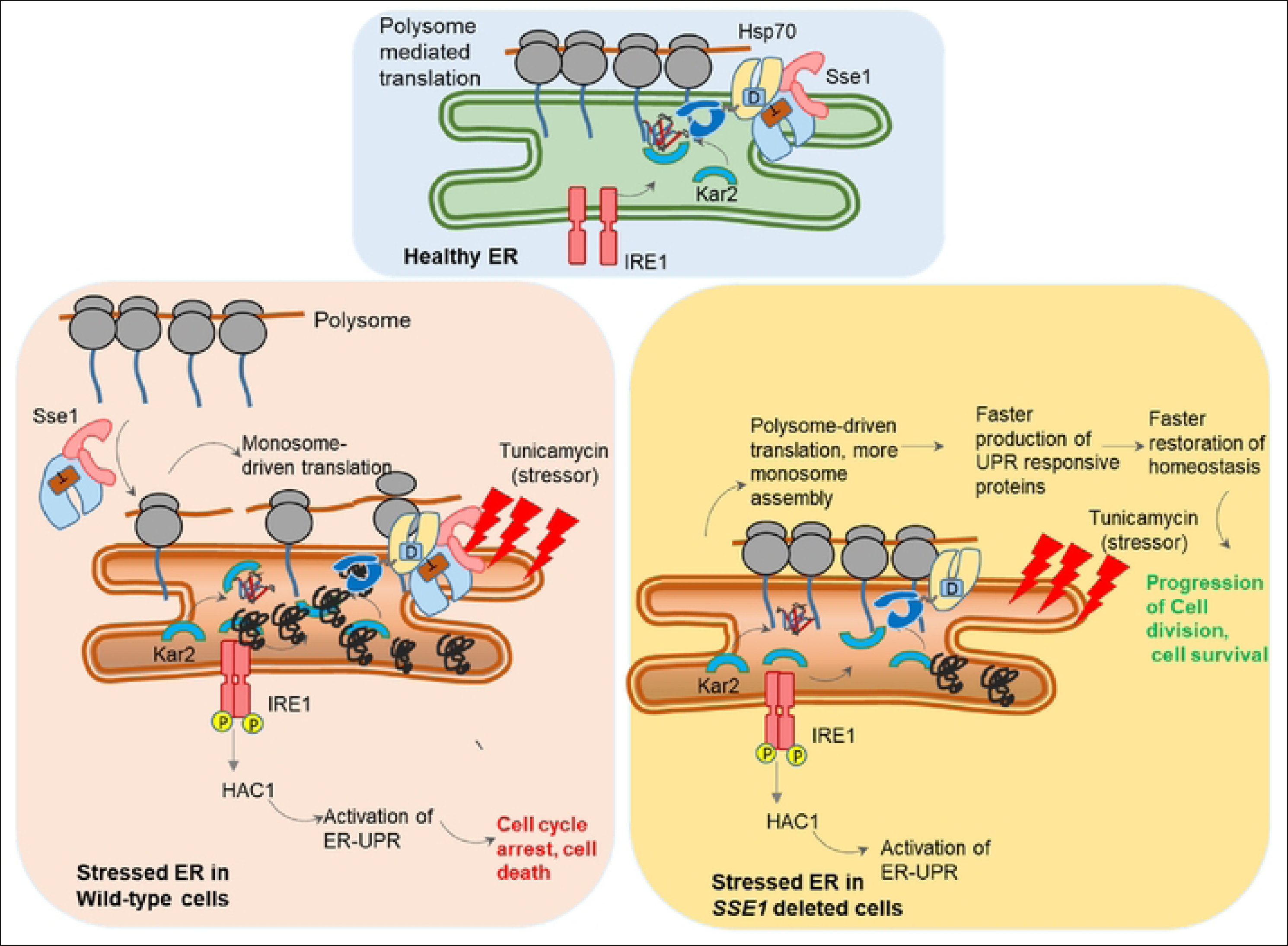
**A schematic summary of the role of Sse1 during ER stress** A schematic model summarizing the role of Sse1 during ER stress. Upper blue box represents physiological condition when there is no stress to ER. The lower left light orange box summarizes the condition of WT cells during Tm-mediated ER stress where polysomes are majorly reorganize to monosomes, and in this process Sse1 plays an important role. There is UPR induction by activation of Ire1-Hac1 pathway which restores the homeostasis upto a tolerable level of stress. The lower right yellow box summarizes the condition of *sse1*Δ cells during ER stress with tunicamycin. Polysome to monosome conversion is inefficient along with more monosome assembly, in absence of Sse1 which leads to faster production of UPR-responsive proteins which restores the homeostasis faster. Additionally, the ER stress induced cell cycle arrest is evaded in *sse1*Δ cells leading to fitness advantage during tunicamycin stress and more cell viability.

It was previously shown that Tm-induced ER stress is known to cause cytokinesis arrest in yeast [27, 28]. As *sse1Δ* strain continued to grow in the presence of Tm-stress, it prompted us to check the cell division status of this strain following ER stress. Importantly, a cell cycle analysis revealed that the cytokinesis arrest, observed in WT cells following long-standing Tm-induced ER stress, is absent in *sse1Δ* strain. The progression of cell division in the *sse1Δ* strain can explain the significantly higher rate of synthesis of new proteins as detected by AHA incorporation by Click-IT reaction following ER stress in contrast to WT cells. Importantly, the reversal to the basal state from the UPR-activated state (post 24 hrs of Tm-treatment) coincided with the reversal of the 3C/4C population to 1C/2C states in cell cycle analysis indicating progression of the cell cycle in the *sse1Δ* strain. Thus, a quicker reversal from the UPR-activated state of the *sse1Δ* strain can be due to cell division and new cell formation. This data nicely corroborated with the unique enrichment of mitotic cell cycle and cell cycle pathways as differentially upregulated pathways exclusively in the transcriptome of Tm-treated *sse1Δ* cells, as described before (Figure 5C). This is important to mention here that we observed differential overexpression of the MAP kinase Slt2 by quantitative mass spectrometry which is critical for yeast cell wall integrity (CWI) [30–32] upon Tm-treatment in both WT and *sse1*Δ cells (Figure 6C and D and Figure S3C, right panel). Importantly, Slt2 was also shown to prevent cell division during Tm-induced ER stress by septin ring mislocalization and cytokinesis block [27, 28]. Additionally, a previous study described that Slt2 interacts with Sse1 and is partially dependent on Sse1 for its cellular activities [33]. Thus, despite overexpression, Slt2 can be inefficient in septin ring mislocalization and cytokinesis block in *sse1*Δ strain leading to cell division progression. Furthermore, upon comparing the proteomics data of both the Tm-treated strains, *sse1*Δ strain showed differential overexpression of Nim1 kinase, Gin4 (Figure 6E), which is a known yeast bud-neck protein that helps in septin collar assembly in the bud neck, promoting cell division [34–38]. Thus, in the presence of an inefficient Slt2, simultaneous overexpression of Gin4 a known factor crucial for septin-collar assembly and bud-neck formation, in *sse1Δ* strain would efficiently counteract the cell division arrest function of Slt2 and promote cell division. Whether Slt2’s inefficient function in arresting cytokinesis or the activities of Gin4 in promoting cell division progression, drives the cell division in *sse1Δ* strain, remains to be understood.

We think that continued protein translation of the *sse1*Δ strain leading to quicker ER-UPR induction and restoration of homeostasis leads to escape from cell division arrest and fitness during Tm-stress (summarized in Figure 8). Taken together, our data show the importance of cytosolic chaperone Sse1 in maintaining ER proteostasis in physiological conditions and during overwhelming ER stress by stressors like tunicamycin. The molecular mechanism of Sse1’s role in controlling cell division arrest during ER stress and how the process is escaped in the absence of the chaperone, remains to be explored in detail.

## Materials and methods

### Yeast strains and associated mutants

We used the yeast (*Saccharomyces cerevisiae)* strain BY4741 (MATa *his3Δ1 leu2Δ0 met15Δ0 ura3Δ0)*, as our wild type and background strain for specific deletion strain. All the *sse1*-double mutants were generated on the commercially available yeast knockout strains from the YKO library employing homologous recombination by using *sse1*-locus-specific *His3MX6* cassette, which was PCR amplified usingpFA6a-His3MX6 plasmid as template and primers containing *sse1*-flanking and *His3MX6* sequence. The other yeast strain which we used is *yMJ003* containing the genotype *MATα his3Δ1 leu2Δ0 met15Δ0 ura3Δ0 LYS+ can1Δ::STE2pr-spHIS5 lyp1Δ::STE3pr-LEU2 cyh2 ura3Δ::UPRE-GFP-TEF2pr-RFP-MET15-URA3*. Apart from this, all the strains that expressed model WT control or stressor proteins were created in the *yMJ003* background as reported previously in detail. Various plasmid purification and yeast transformation were carried out using standard laboratory protocols.

### Yeast culture and growth assay

The strains from the yeast knockout library of the BY4741 background were grown in the YEPD (1% Yeast extract, 2% Peptone and 2% Dextrose) for overnight to get a saturated culture and the next day a secondary culture was inoculated at 0.1 OD_600_. To perform the drop dilution assay, secondary cultures of specific yeast strains were grown till the mid-log phase (0.4-0.6 of OD_600_), and serially diluted and spotted on different media plates as mentioned in each experiment. For keeping plasmids, synthetic media with particular auxotrophic selection were used. For the ER-MBP and ER-DMMBP strains made in the YMJ003 background, poorly fermentable YPR (1% Yeast extract, 2% Peptone and 2% Raffinose) media was used as control. 1% galactose (final) was used as the inducer for the expression of MBP or DMMBP proteins from the Gal1 promoter in the YPR media. Liquid growth assay was done using the Bioscreen instrument (Clover Biotech).

### Polysome Profiling

Specific yeast strains were reinoculated in the secondary cultures at 0.1 OD_600_ from the primary cultures and allowed to grow till 0.4-0.6 OD_600_. After that, the desired treatments were given according to the experimental setup. After the treatment was done, cycloheximide (50µg/ml) was added to the media and incubated on ice for 5 minutes. Then the cells were harvested at 2500 RCF for 10 minutes at 4°C. Then the cells were resuspended in lysis buffer (50 mM Tris pH - 7.5, 150 mM NaCl, 30 mM MgCl_2_, 50 μg/ml Cycloheximide) and lysed with the bead beater with the settings 15 seconds On and 30 seconds Off for 10 cycles. In between the cycles, the samples were incubated on ice. After this, the cells were centrifuged at 4000 RPM for 5 minutes at 4°C and then the supernatants were collected in a fresh tube and spun at 9200 RPM for 10 minutes at 4°C. After this, the total RNA content of each strain was normalised by taking absorbance at 260 nm in Nanodrop to proceed with ultracentrifugation. From each, the strain equal amount of RNA (Here 10 absorbance units at 260 nm) should be loaded to the 7 - 47% continuous sucrose gradient (50 mM Tris Acetate pH - 7.5, 250 mM Sodium Acetate, 5 mM MgCl_2_, 1 mM DTT and 7, 17, 27, 37, and 47% sucrose) and then centrifuged in Beckman Coulter SW32Ti rotor at 165000 RCF for 4 hours at 4°C. After the ultracentrifugation, all the samples were fractionated using the ISCO gradient fractionator with the UV detector sensitivity set to 0.5 or 1.0 (the sensitivity should be optimized according to the initial load and the peaks in the profile).

### CLICK Chemistry

We used CLICK chemistry to assess the new protein synthesis rate under normal and stressed conditions in various yeast strains. The specified strains required by the experimental setup were reinoculated in secondary cultures at 0.1 OD_600_ from the primary cultures and allowed to grow till 0.4-0.6 OD_600_ in synthetic complete media. After that cells were collected at 5000 RPM for 5 minutes, washed with sterile water and then centrifuged at 5000 RPM for 5 minutes. Then resuspend the cells in SD-AHA (Standard media where Methionine is replaced with L-Azidohomoalanine) and treated with tunicamycin as per the experimental setup. Then the cells were harvested at 8000 RPM for 5 minutes, washed with sterile water and then centrifuged at 8000 RPM for 5 minutes following which the supernatant was discarded. Then to permeabilize the cells they were resuspended in 53% (v/v) molecular grade absolute ethanol in 1X PBS and incubated in an incubator shaker set at 14.8°C with 200 RPM shaking for 40 minutes [2]. After that the cells were collected by centrifugation at 8000 RPM and the supernatant was discarded. The cells were then incubated with 1 ml of CLICK reaction cocktail (2M Tris-pH8.5, 50mM Copper Sulphate, 1µg/ml Alexa-Fluor 488 alkyne, and 0.5M Ascorbic acid) for 30 minutes at room temperature. Following that the cells were collected at 8000 RPM for 5 minutes washed with 1 ml of 1X PBS and then resuspended in 300µl of 1X PBS and proceeded for Flow Cytometry as well as Confocal Microscopy analysis.

### Statistical Analysis

Descriptive statistics for all the measurements that are used for plotting the graphs are expressed as mean ± standard error of the mean (SEM). For calculating the statistical significance, we performed an unpaired T-test and one-way ANOVA. If one-way ANOVA yielded a significant difference, then we performed Tukey’s test as a post hoc analysis for pairwise comparisons. We set the significance threshold α at 0.05 for all statistical testing that was performed assuming equal variance and the significances were calculated at the two-tailed level. In cases where multiple tests were performed Bonferroni Correction was used to account for the family-wise error rates. The calculated p-values were signified as stars in the plots according to the following manner: ‘ns’ meaning P > 0.05, ‘*’ meaning P ≤ 0.05, ‘**’ meaning P ≤ 0.01, ‘***’ meaning P ≤ 0.001, and ‘****’ meaning P ≤ 0.0001 respectively. For mass spectrometry data analysis, the fold change analysis was done on the raw expression values; following which the fold change values were transformed to a logarithmic scale having base 2 and termed as ‘Log_2_-Fold Change’. The statistical significance of the expression values was calculated using unpaired T-tests assuming equal variance on the Log_2_ transformed expression values. The resulting p-values were transformed to a negative logarithmic scale having base 10 and termed as ‘-Log_10_(P)’. Finally, the volcano plots were created using the ‘Log_2_-Fold Change’ values on the X-axis and the ‘-Log_10_(P)’ values on the Y-axis.

Other methodologies have been described in the supplementary methods.

## Acknowledgements

KM acknowledges the funding support from the Science and Engineering Research Board (SERB), Government of India, for Core Research Grant (SERB/CRG/2019/006281) and SNU core funding. MPJ acknowledges the SNU PhD fellowship and ICMR SRF Grant (2020-4242/CMB-BMS). MJ and KM acknowledge the SNU DST-FIST grant [SR/FST/LS-1/2017/59(c)] for the confocal microscopy facility. We thank Praveen Singh for his help with mass spectrometry data analysis. We thank Manisha Kochar for helping with FACS (BD-LSRII) measurement for cell cycle analysis. We thank Akanksha Sharma for helping with the yeast liquid growth assay using the Bioscreen instrument. We thank Dr Neelesh Dahanukar for his help in various statistical analyses. We thank Dr Kausik Chakraborty for sharing the YMJ003 yeast strain and allowing us to use the BD-LSRII flow cytometer and Bioscreen instrument for yeast liquid growth assay at CSIR-IGIB.

## Author Contribution

The work was conceived by KM. All yeast strains were generated by MPJ and VK. Most yeast experiments were done by MPJ and some initial experiments were done by VK.. Flow cytometry experiments and cell-cycle analysis, biochemical assays and cell death assays, microscopy and data analysis were performed by MPJ. RNA sequencing was run by AG and VK and data were analysed by AG and MPJ. LS ran the proteomics samples and data were analysed by MPJ. KM supervised the work, analysed the data and wrote the manuscript.

---

**Figure S1.**
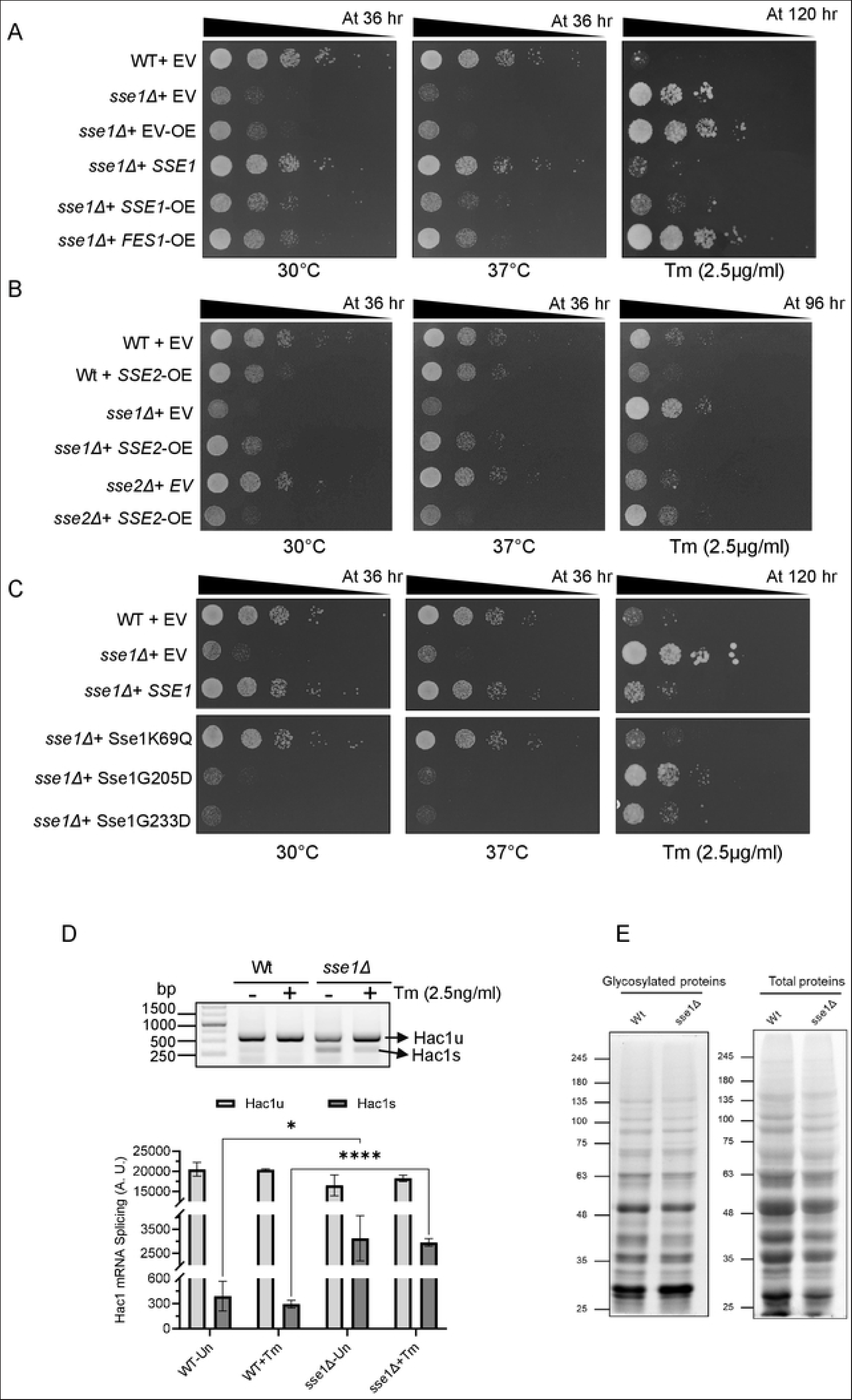

**Figure S2.**
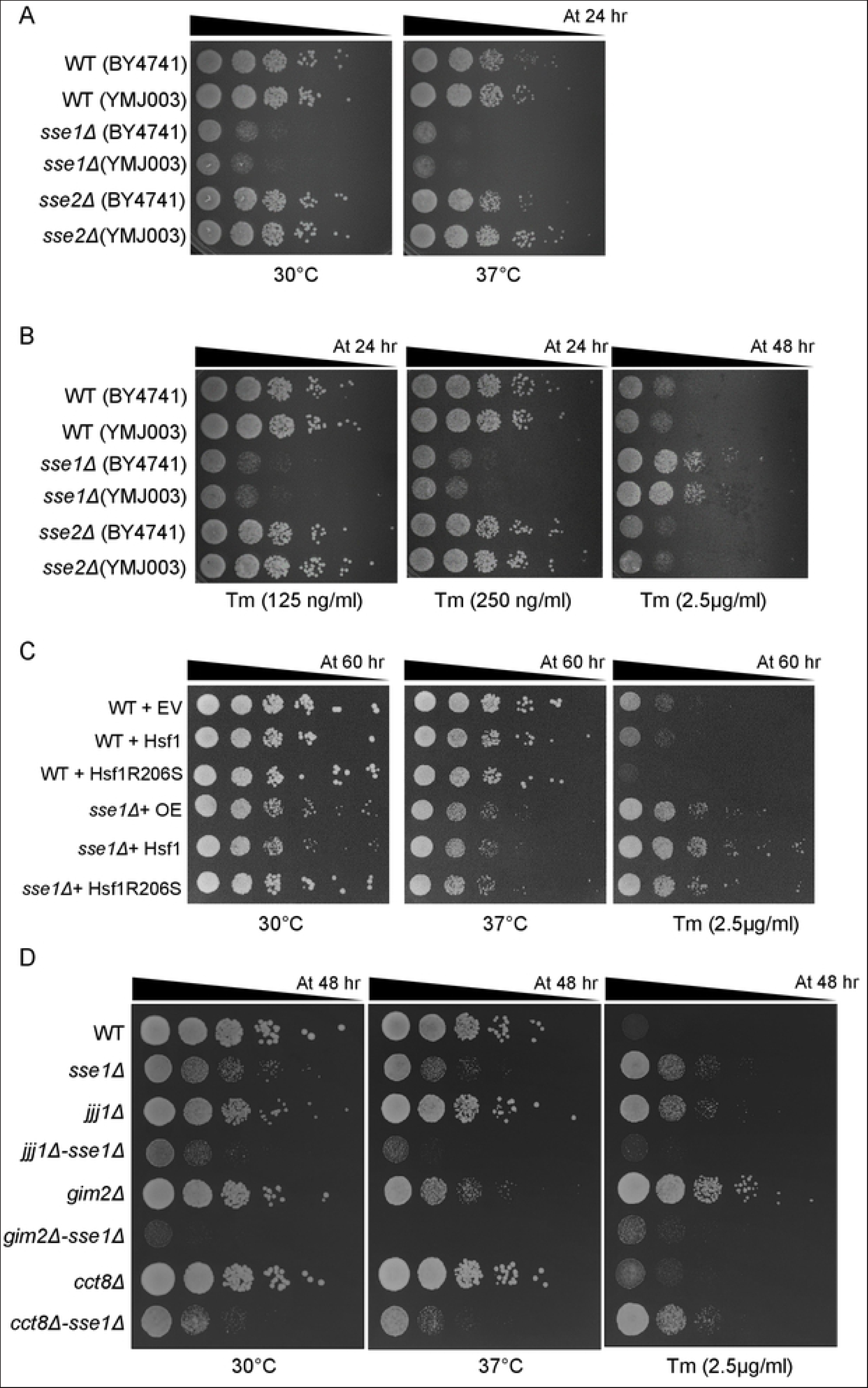

**Figure S3.**
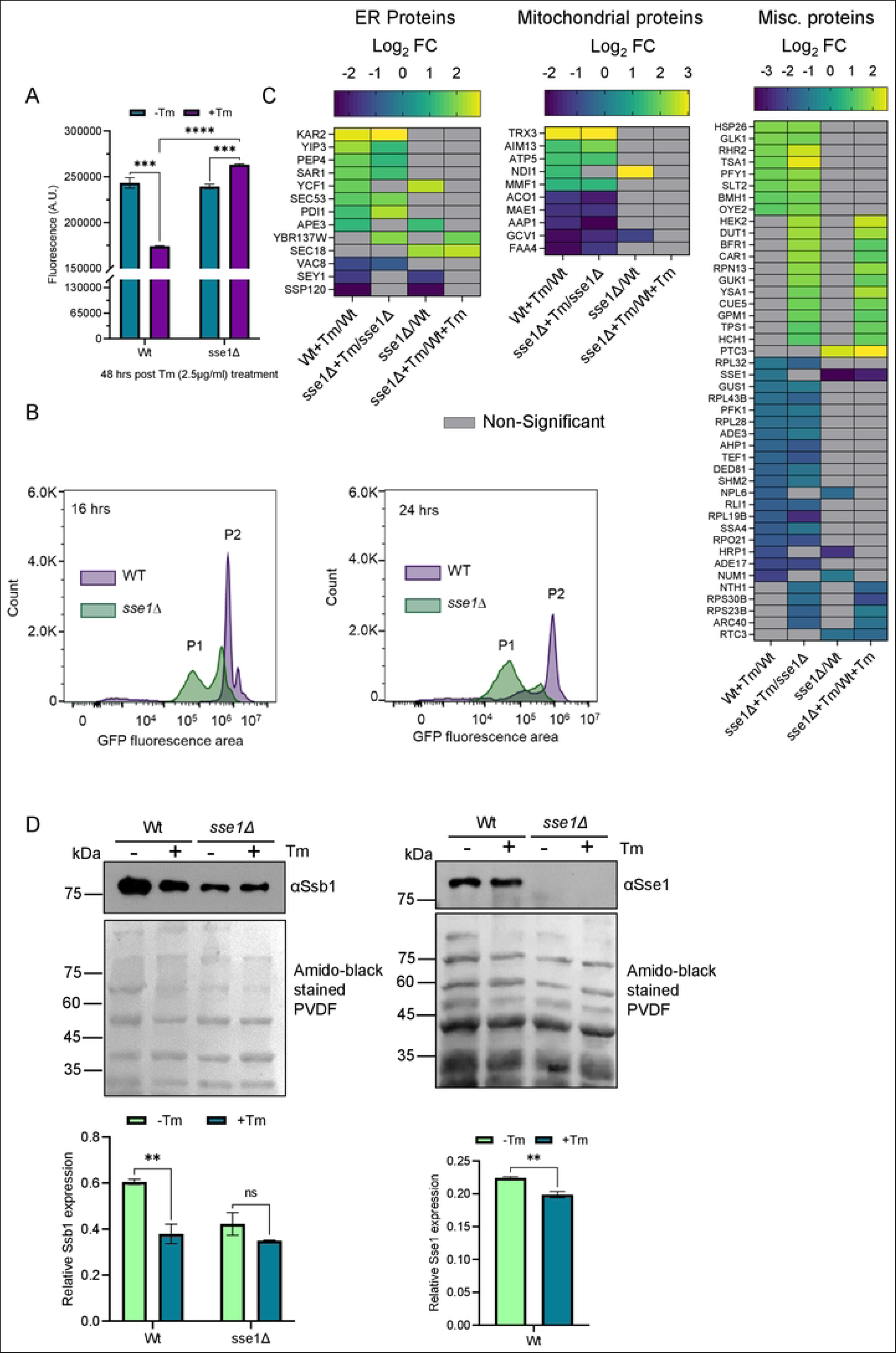

**Figure S4.**
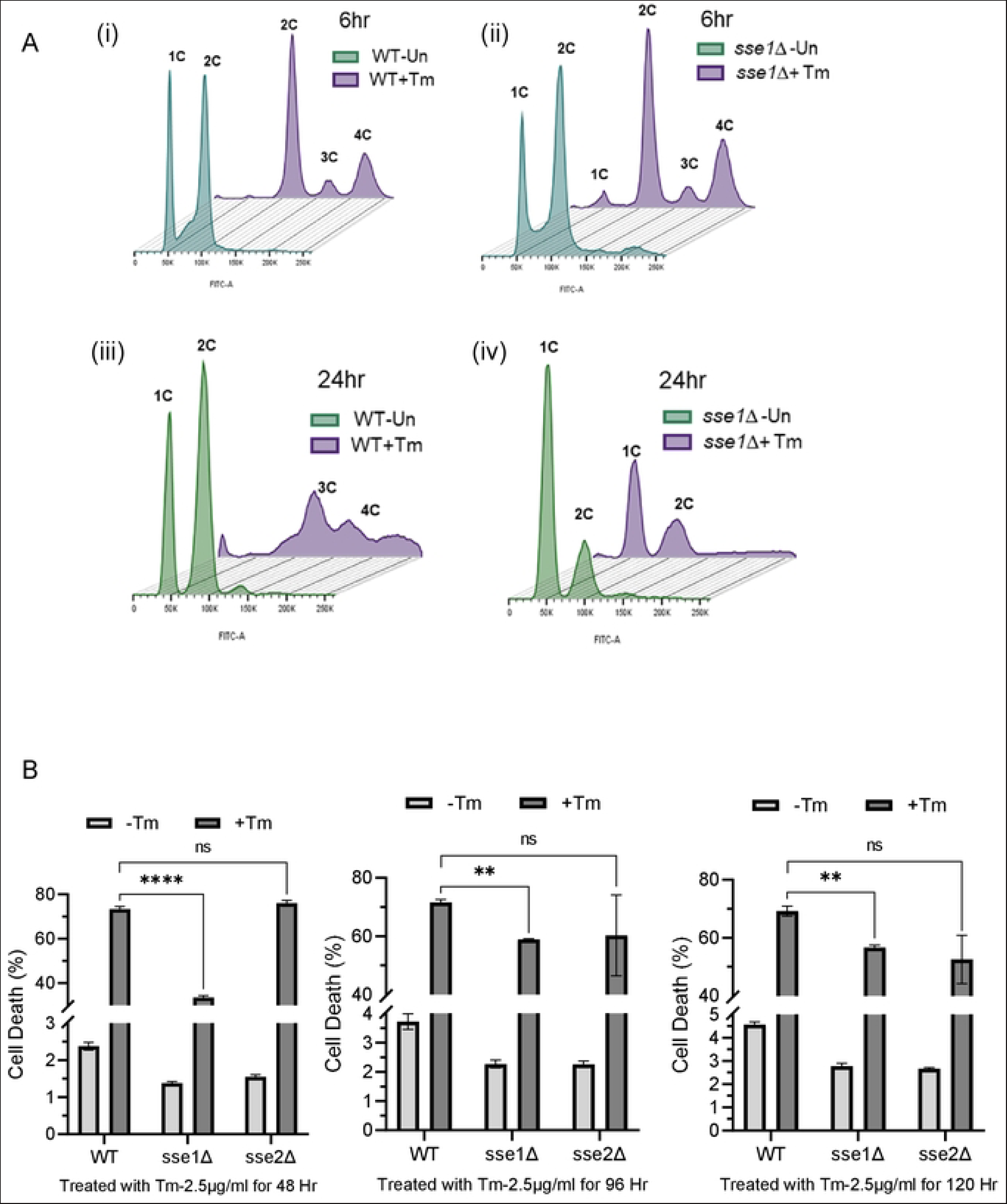

